# ETS TRANSCRIPTION FACTOR POINTED CONTROLS GERMLINE SURVIVAL IN *DROSOPHILA*

**DOI:** 10.1101/2024.08.15.608137

**Authors:** Alicia E. Rosales-Nieves, Miriam Marín-Menguiano, Lourdes López-Onieva, Juan Garrido-Maraver, Acaimo González-Reyes

## Abstract

Proper gonad development is a pre-requisite for gametogenesis and reproduction. During female gonad formation in *Drosophila*, the EGF receptor (EGFR) signalling pathway ensures the correct number of primordial germ cells (PGCs) populate the larval gonad. We study the gene *pointed (pnt)*, which acts downstream of the EGFR receptor and belongs to the ETS transcription factor family, with a previously unknown function in gonadogenesis. We report that *pnt* is expressed in female larval gonads and later in the adult ovarian germline niche and that it is required to sustain proper gametogenesis. Loss of *pnt* function in female larval gonads, similar to the EGFR, induced PGC overproliferation. Conversely, we isolated a novel loss-of-function allele, *pnt^aga^*, which resulted in agametic gonads and ovaries. While *pnt^aga^* embryos developed gonads containing a normal complement of PGCs, these are subsequently lost by apoptosis during late larval and pupal stages. Molecular characterization of *pnt^aga^* revealed reduced expression levels of the different *pnt* isoforms, unveiling a complex autoregulatory network involving the three Pnt proteins. We propose that germ line survival in *Drosophila* gonads requires a precise tuning of EGFR signalling to ensure the appropriate transcriptional activation of its target *pnt*.

## 1. Introduction

Understanding how tissue specification and morphogenesis are controlled is a central question in biology. During animal development, multiple processes are tightly orchestrated in time and space to facilitate specific tissue differentiation and organ formation. *Drosophila* oogenesis serves as an elegant model system in this regard, allowing for extensive analysis of cell behavior and organ morphogenesis in a genetically tractable and experimentally flexible context. Here, we report the isolation of a new mutation in the *pointed* (*pnt*) gene that halts germline development and provides key insights into morphogenesis.

Pointed belongs to the ETS-domain family, one of the largest groups of transcription factors described so far (Wang et al., 2023). This family of proteins, unique to metazoans, encompasses over 30 proteins from humans to *Drosophila*. ETS (E26 Transformation Specific) transcription factors are implicated in critical developmental processes as well as in cancer progression, and they have been known to regulate a myriad of cellular and viral genes (Dittmer & Nordheim, 1998; Wang et al., 2023). They are characterized by the presence of an evolutionary conserved DNA-binding domain (the ETS domain) and act in cooperation with a variety of structurally unrelated transcription factors and co-factors (Sharrocks, 2001). Most ETS proteins are nuclear targets of Ras-MAP kinase signaling cascades and regulate cell proliferation, differentiation and apoptosis by activating genes encoding growth factor receptors and integrin families, among others (reviewed in Oikawa & Yamada, 2003; Verger & Duterque-Coquillaud, 2002). Furthermore, studies in model organisms such as *C. elegans* and *Drosophila* have uncovered the critical role of these transcription factors in a variety of developmental processes such as oogenesis, metamorphosis, neurogenesis, eye development and myogenesis (Hsu & Schulz, 2000).

The *Drosophila pointed* locus gives rise to four transcription variants: *pnt-RB, -RC, -RD* and *-RE*. Since *pnt-RC* and *pnt-RE* only differ in their 3’UTR, the locus gives rise to three different protein isoforms: Pnt-PB, Pnt-PD and Pnt-PC/E (hereafter we will refer to the three products as long, intermediate and short; Fig. 1A, B). The three Pnt proteins share an ETS domain, common to all members of the ETS family (Karim et al., 1990). However, only Pnt^long^ and Pnt^intermediate^ incorporate a second conserved homology domain named POINTED (PNT), also known as SAM-PNT (for <u>S</u>terile <u>A</u>lpha <u>M</u>otif), which becomes phosphorylated upon activation of the upstream Ras-MAPK pathway (Fig. 1B) (Klambt, 1993; Morimoto et al., 1996; O’Neill et al., 1994). Further allelic characterization of *pnt^long^* promoter described its function during eye and wing development and identified two sequence domains with opposite effect on transcription, one *trans*-activating and another *trans*-repressing transcription of *pnt^long^* (Scholz et al., 1993). Contrariwise, Pnt^shor*t*^ is constitutively active because it lacks the phosphorylation domain. The original model suggested that Pnt^long^, which responds to the EGFR/Ras/MAPK pathway, may turn on the *pnt^short^* promoter, thus allowing a prolonged activation signal in the target cell (O’Neill et al., 1994). Both isoforms typically work in concert with their transcriptional partner, Yan, which serves as a negative regulator (Boisclair Lachance et al., 2014; O’Neill et al., 1994; Vivekanand et al., 2004; Webber et al., 2018). Specifically in the context of eye development, Yan operates in opposition to the Ras signalling function (Brunner et al., 1994; O’Neill et al., 1994). It was originally proposed that *yan*, which itself encodes another ETS-related transcription factor, competed with Pnt for binding targets (O’Neill et al., 1994). More recently, however, it has been suggested that Pnt promotes the recruitment and stabilization of Yan and co-repressor Groucho. Upon activation of EGFR signaling, disassembly of the Yan, Pnt and Gro complex enables RNA polII transcription elongation (Webber et al., 2018). The highly conserved ETS domain binds DNA over a range of 12 to 15 bp, exhibiting sequence preference for around 9 bp with a central invariable core: 5’-GGA(A/T)-3’ (Hollenhorst et al., 2011). The transcriptional activity of the *Drosophila* Pnt^long^, and its mammalian orthologs ETS1 and ETS2, depends on mitogen-activated protein (MAP) kinase phosphorylation. The PNT domain works as a docking sequence to enhance phosphorylation of an immediately adjacent threonine by, in the case of *Drosophila* Pnt^long^, the MAP kinase Rolled (Brunner et al., 1994).

**Figure 1:**
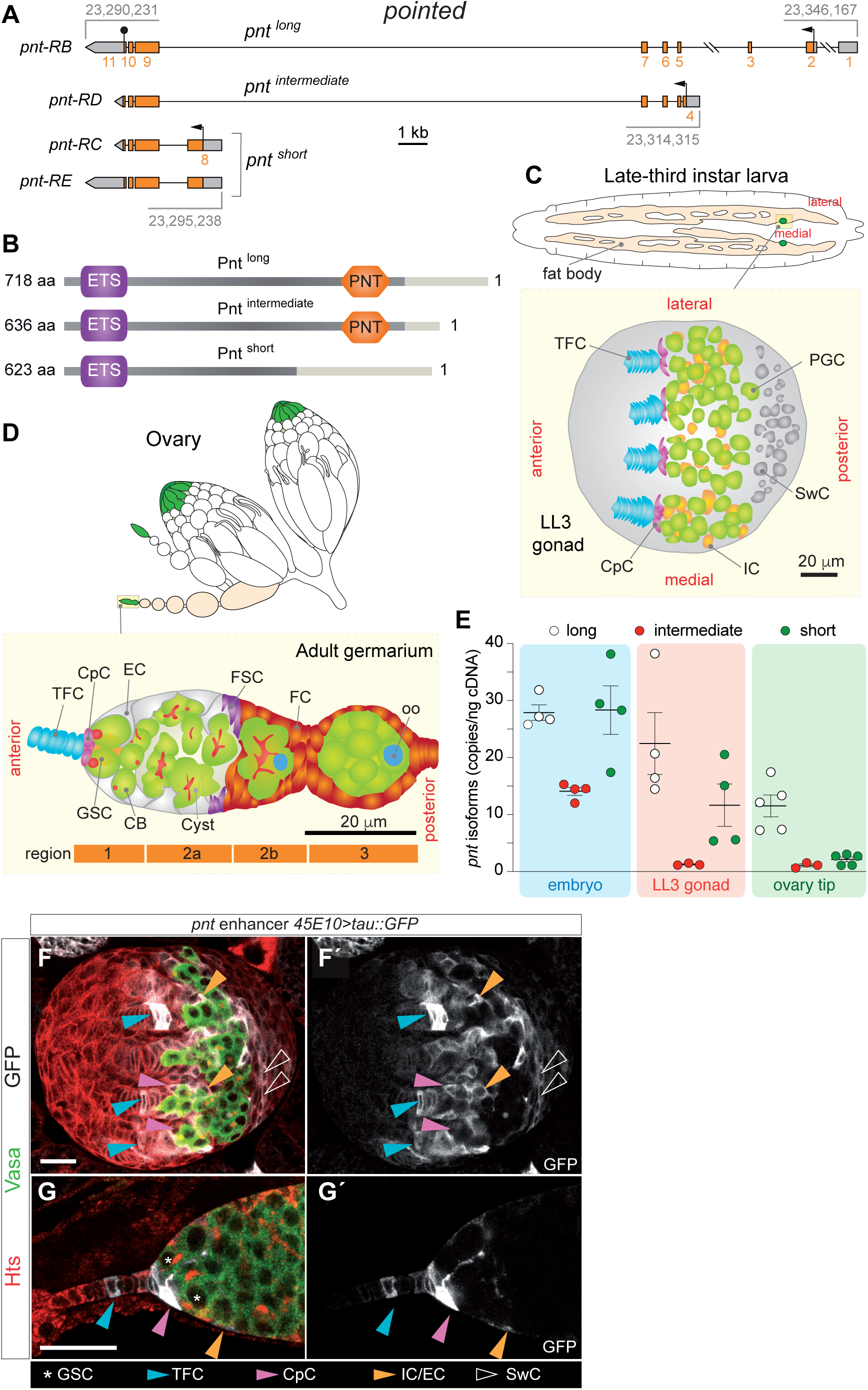
Pointed is expressed in female larval gonads and in the ovarian GSC niche. **(A)** Scheme of the *pointed* locus highlighting the four predicted isoforms. Arrows designate transcriptional start sites. The stop codon is indicated by the black sphere. UTRs (untranslated regions) are shown in grey. The three different ORFs (open reading frames) are coloured in orange and the exons are numbered in orange. Numbers in grey refer to genomic coordinates according to FlyBase (release FB2024_02). **(B)** Scheme of the three Pointed protein isoforms. The PNT (Pointed) and the ETS (E26-Transformation Specific) domains are highlighted. Light grey boxes represent isoform-specific domains. **(C)** Drawing of a female LL3 larva showing the fat body and the medial position of the embedded gonads. Drawing of an LL3 gonad revealing the distribution of relevant cell types. **(D)** Drawing of a pair of ovaries to highlight the organization of ovarioles and the localization of germaria at their tip. Drawing of a germarium to show the distribution of relevant cell types. **(E)** Detection of the *pnt-RB* (long), *-RD* (intermediate) and *-RC/E* (short) mRNAs using droplet-digital PCR in control embryos, LL3 gonads and ovary tips containing germaria and few early egg chambers. The primer pairs specific for each of the three isoforms map to exons 2 and 5 (long), 4 and 5 (intermediate) and 8 and 9 (short). Measurements correspond to at least three biological replicates and to one or two technical replicates. **(F, G)** Pattern of expression of the *pointed* 45E10 enhancer in LL3 gonads (F) and in germaria (G). 45E10-driven expression of the Tau::GFP reporter is shown in white, Hts in red and Vasa in green. Scale bars: 20 μm. In this and the rest of the figures, abbreviations used are the following: TFC: terminal filament cell; CpC: cap cell; EC: escort cell; GSC: germline stem cell; PGC: primordial germ cell; CB: cystoblast; FSC: follicle stem cell; FC: follicle cell; oo: oocyte; IC: intermingled somatic cell; SwC: Swarm cell. Related to Figure S1.

In *Drosophila* females, the Germline Stem Cell (GSC) niche begins to shape at larval stages when the gonad consists of a growing, round mass of somatic connective tissue and primordial germ cells (PGCs), surrounded by fat body located posteriorly (Fig. 1C). In third larval instar, the anterior of the gonad contains the precursors of the terminal filament cells (TFCs) and cap cells (CpCs); the medial-lateral axis houses the PGCs and somatic interstitial cells called intermingled cells (ICs); and the most posterior region accommodates the swarm cells (SwCs; Fig. 1C). The TFCs organize in stacks following a morphogenetic wave that moves medial to lateral and configures the final organization of the terminal filaments (TFs). The GSC niche is complete by early pupa, when TF formation is concluded and CpCs are incorporated at the base of the TFs (Godt & Laski, 1995; Sahut-Barnola et al., 1995; Yatsenko & Shcherbata, 2018). During larval stages, PGCs proliferate, increasing their number from ∼12 in the embryo to about 100 undifferentiated germline cells by mid-third instar larvae (ML3). PGCs duplicate numbers every 24h during first and second instar, then the rate decreases during the following 24h (Gilboa & Lehmann, 2006). PGCs are prevented from differentiating until ML3, when enough PGCs exist to populate all the niches. The PGCs in contact with the CpCs cells at the base of the TFs will become Germline Stem Cells (GSCs), while the PGCs that end up away from the niche will enter differentiation. After ovariole individualization during pupal stages, the adult ovary is formed by 16-18 ovarioles where mature eggs are produced. At the anterior end of each ovariole, a conical structure called germarium harbors the GSC niche, a tightly regulated system of 2-4 stem cells and 3 different somatic cell types - CpCs, in direct contact with the GSCs; TFCs, at the very tip, forming the TF; and anterior escort cells (ECs) (Fig. 1D). The GSC niche provides the signaling required for GSC maintenance and proliferation, allowing the generation of mature eggs throughout the lifetime of the fly.

The first characterizations of the *pnt* locus in *Drosophila* reported that it is required for proper central nervous system development (CNS) in embryos and for the morphogenesis of compound eyes. Embryonic *pnt* loss-of-function phenotypes include fusion of the CNS commissures and abnormal glial-neuron interactions (Klambt, 1993; Scholz et al., 1993). In the adult, non-lethal allelic combinations of *pnt* present a rough eye phenotype, which could be indicative of defects in cell fate acquisition (Scholz et al., 1993). In fact, Pnt is a target of Ras signaling controlling photoreceptor determination and it is activated *in vitro* by MAP kinase phosphorylation (Brunner et al., 1994; O’Neill et al., 1994). In this study, we investigate the role of *pnt* in the context of *Drosophila* gametogenesis. We characterize a new mutation in the *pointed* locus that is homozygous viable but causes sterility, as the reproductive system of mutant adults is agametic. Our results demonstrate the importance of the correct transcription of the locus for proper gonadogenesis and hence gametogenesis and reproduction.

## 2. Results

### Pointed is expressed in the somatic cells of the female larval gonad and ovarian GSC niche

*pnt* spans ≈56 kb and encodes four different transcriptional variants. The transcription start sites (TSS) of *pnt^long^* and *pnt^short^*, are separated by ≈50 kb of mostly intronic sequence. The TSS of *pnt^intermediate^* falls within this interval (Fig. 1A). Considering the essential function of the EGFR pathway for proper female gonadogenesis and that *pnt* is a downstream target of the pathway (Gilboa & Lehmann, 2006; Morimoto et al., 1996), we studied a potential role for *pointed* in regulating GSC proliferation and/or niche formation throughout development. We used droplet digital PCR (ddPCR) to measure expression levels of the long, intermediate and short isoforms in embryos, female larval gonads and during early oogenesis in the adult. All three isoforms were found in embryos, consistent with the known roles of *pnt* in embryonic central nervous system development (Klaes et al., 1994; Klambt, 1993). In contrast, female late third instar larval (LL3; 5 days after egg laying) gonads only expressed *pnt* short and long, while in ovary tips (germaria plus few, early egg chambers) we could only detect consistent levels of *pnt* long (Fig. 1E).

To analyze further *pointed* expression in LL3 female gonads and throughout adult oogenesis, we utilized a number of lines expressing the Gal4 trans-activator under the control of different *pnt* enhancers and monitored their expression using a *UASp-Tau::GFP* reporter line. In all ten lines analyzed, we detected *pointed* expression in somatic cells of the larval gonad including TFCs, CpCs and SwC. In the adult ovary, *pnt-Gal4*-driven Tau::GFP was consistently observed in TFCs, CpCs and in anterior ECs in the germarium (Fig. 1F, G; Fig. S1). In addition, and as described before (Morimoto et al., 1996, Lee & Montell, 1997, and Meignin et al., 2007), we could also detect expression of *pnt* in follicle cells at the posterior of the egg chamber from stage 6, as shown for the *Gal4* line NP5370, inserted within the *pnt* locus (Fig. S1). Thus, *pnt* expression seems to be restricted to the somatic component of the female gonad and adult ovary.

### *pointed* restricts PGC numbers in the female gonad and ensures cyst encapsulation in the germarium

*pnt* null alleles are embryonic lethal (Scholz et al., 1993). Thus, studying *pnt* function during gonad formation requires either genetic tools to manipulate gene function in groups of cells or the use of hypomorphic mutations that give rise to viable adults. To evaluate the role of *pnt* in early oogenesis, first we used the UAS/Gal4 system to induce RNAi-mediated gene silencing. We examined the effect of lowering *pnt* expression in the larval gonad and the adult germarium by expressing either of two RNAi lines (KK100473 and JF02227) under the control of *c587-Gal4* (*c587>pnt*^RNAi^), which is expressed in the somatic cells of the gonad (Zhu & Xie, 2003) and germarium (Kai & Spradling, 2003) (Fig. S2). We observed a significant increase in PGC numbers for either of the RNAi lines compared to controls in female larval gonads (Fig. 2A-C; similar results were obtained for the *traffic jam-Gal4* line, also expressed in the somatic cells of the gonad; Fig. S2). Of interest, preventing EGFR pathway activation in the somatic cells of the female larval gonad gave rise to excess of PGCs (Gilboa & Lehmann, 2006; Matsuoka et al., 2013). Our results thus suggest that *pnt* acts downstream of the EGFR cascade in the somatic cells of the female gonad to control proliferation and survival of PGCs. The expression of the JF02227 RNAi construct with the *c587-Gal4* driver in adult females gave rise to germaria larger than controls containing more germline cells, in agreement with a previous report (Liu et al., 2010; Fig. 2D-E). In addition, we also observed egg-chamber fusions (Fig. 2E), a novel phenotype suggesting that *pnt* is involved in somatic cell fate specification, proliferation and/or migration in the germarium. Since the RNAi lines used in these experiments are designed to target regions common to all the *pnt* isoforms and since the *c587-* and *traffic jam-Gal4* lines are expressed in somatic cells (Fig. S2A, C-D), we conclude from these results that reduction of *pnt* function in the somatic component of female gonads and ovaries controls germline proliferation and gametogenesis.

**Figure 2:**
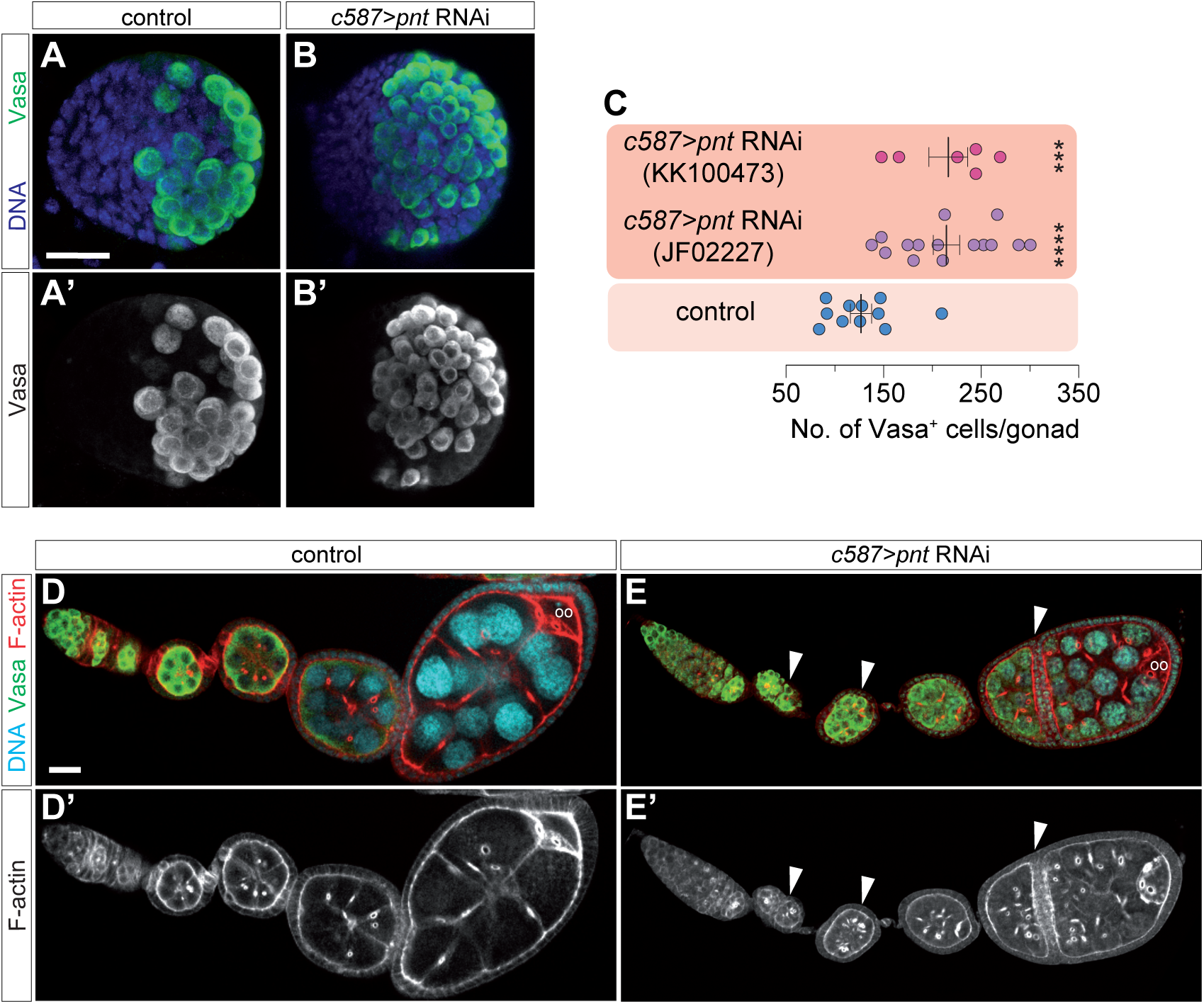
*pointed* restricts PGC numbers in the gonad and ensures proper cyst encapsulation in the germarium. **(A, A’)** Control LL3 gonad stained with anti-Vasa (green) to show PGCs and with a DNA dye (blue) to mark chromatin. **(B, B’)** LL3 gonad from a *c587>pntRNAi^JF02227^* female larva showing the excess of PGCs found in *pnt* loss-of-function gonads. **(C)** Quantification of the number of Vasa+ cells in control, *c587>pntRNAi^JF02227^* and *c587>pntRNAi^KK100473^* LL3 gonads. The arithmetic mean and the SEM are shown for each of the genotypes. **(D, D’)** Ovariole from a control female stained to visualize F-actin (red), the germ line (anti-Vasa; green) and with a DNA dye (blue). Developing egg chambers are formed in the germarium and each contains a 16-cell germline cyst surrounded by a monolayered follicular epithelium. The oocyte is always found at the posterior of the cyst. **(E, E’)** Ovariole from a *c587>pntRNAi^JF02227^* female stained as in (D). White arrowheads point to fused egg chambers containing more than one oocyte and 15 nurse cells. ***=p<0.0005; ****=p<0.0001. Scale bar: 20 μm. Related to Figure S2.

### *pnt^aga^*, a novel mutation that induces germline cell death

We have isolated and characterized a spontaneous mutation in the *pnt* locus that we named *pnt agametic* (*pnt^aga^*). Contrary to the above results, adult *pnt^aga^* homozygous males and females were viable but displayed a highly penetrant agametic phenotype that renders them sterile. This phenotype was not a consequence of abnormal embryonic gonad development, since gonads from control and *pnt^aga^* embryos were indistinguishable (Figs. 3I and S4). In contrast, the phenotypic characterization of ML3 gonads from female *pnt^aga^* larvae (4-5 days after egg laying) already showed significant differences with controls (Fig. 3A). Thus, compared to controls (average 98.22 ± 5 *n* = 9), the mutant PGC pool had an aberrant and disorganized morphology and the number of mutant PGCs at ML3 was considerably reduced (average 55.5 ± 4.35 *n* = 10; Fig. 3A, B, I). To analyse the consequences for gonad maturation of the effects observed in larval tissues, we stained female control and *pnt^aga^* white pupal gonads (∼24 hours later in development) to find that mutant gonads contained greatly reduced PGC numbers compared to controls (controls: average 206.87 ± 8.7 *n* = 8; *pnt^aga^*: average 34± 3.8 *n* = 13). In addition, the few remaining mutant germ cells appeared aberrant in shape and larger in size (Fig. 3C, D, I). Finally, homozygous mutant adult ovaries were considerably smaller than controls and were completely deprived of germ cells including GSCs, cystoblats and cysts. This phenotype was fully penetrant, as 100% of adult ovarioles were negative for the germline marker Vasa (Fig. 3E-H, I). We also used an anti-Engrailed antibody to establish that mutant niches, while depleted of germline cells, seemed to form properly and to contain a normal pool of TFCs and CpCs (Fig. 3G, H). This further confirmed that the reduced ovaries of agametic females maintain some characteristics of normal ovarian niches (Margolis & Spradling, 1995).

**Figure 3:**
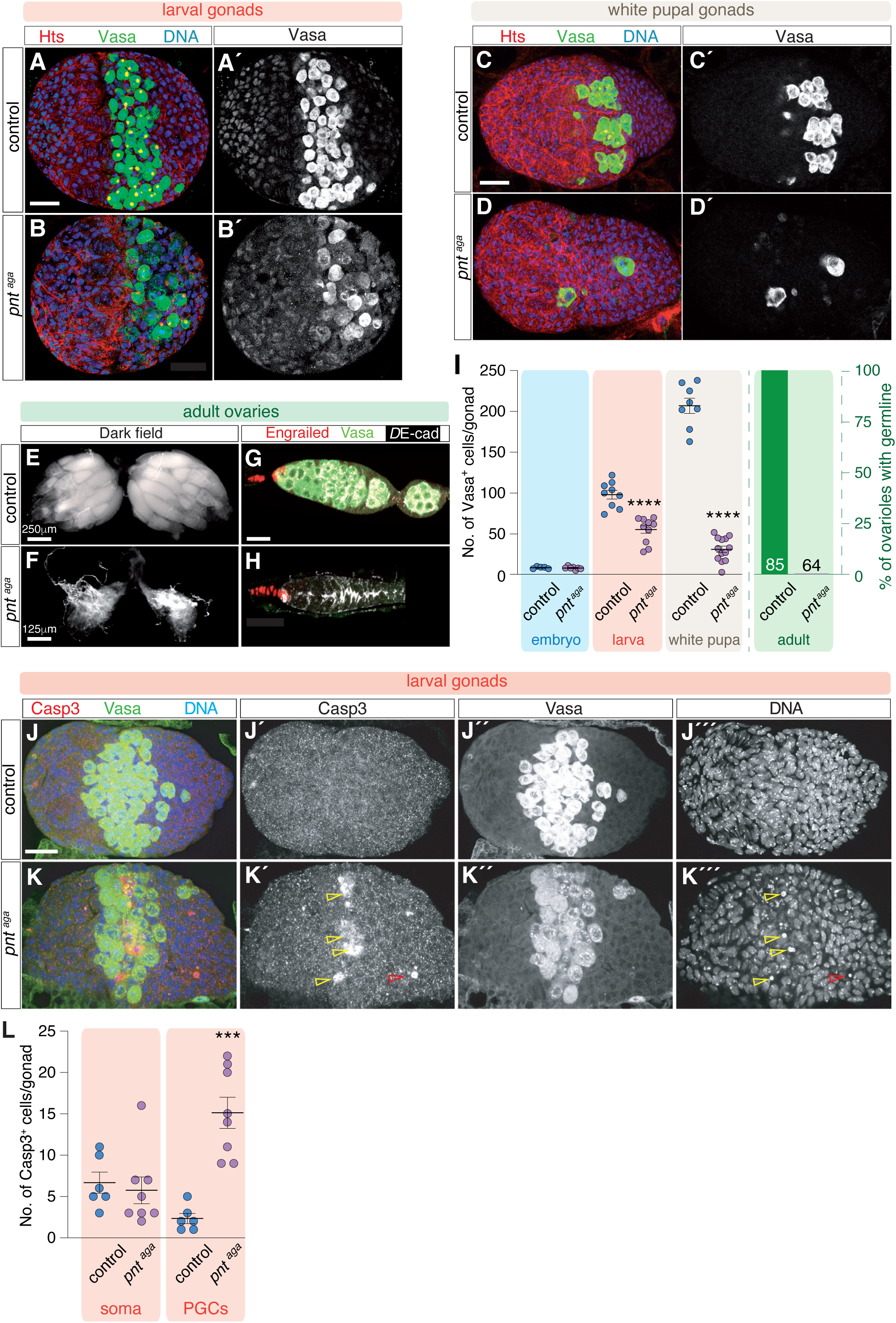
*pnt^aga^* regulates germline survival. **(A)** Control and **(B)** *pnt^aga^* ML3 gonads stained with anti-Hts (red; to visualise cell outlines and germline spectrosomes and fusomes), anti-Vasa (green; to label the germ line) and with a DNA dye (blue; to mark chromatin). **(C)** Control and **(D)** *pnt^aga^* white pupa gonads stained as in (A). Note the progressive decline in germline cells in the mutant condition. **(E, F)** Darkfield images of control and *pnt^aga^* ovaries. **(G, H)** Control and *pnt^aga^* germaria stained with anti-Engrailed (red; to label TFCs and CpCs), anti-Vasa (green) and *D*E-cad (white). Note that *pnt^aga^* ovaries and germarium are devoid of germline cells. **(I)** Quantification of the number of Vasa+ cells in control and *pnt^aga^* embryos, ML3 gonads and white pupa gonads. Quantification of the percentage of ovarioles with germline in control and mutant adult ovaries. Numbers in columns denote sample size (n). The arithmetic mean and the SEM are shown for each of the genotypes. **(J)** Control and **(K)** *pnt^aga^* LL3 gonads stained with anti-Caspase 3 (red; to visualise apoptotic cells), anti-Vasa (green) and with a DNA dye (blue). **(L)** Quantification of the number of Casp3+ cells in somatic or germline cells in control and *pnt^aga^* gonads. ***=p<0.0005; ****=p<0.0001. Scale bars: 20 μm unless otherwise noted. Related to Figures S3 and S4.

To test whether the agametic phenotype observed in mutant ovaries is due to PGC death concomitant with niche formation, we stained ML3 gonads with anti-Caspase 3, a broadly used marker for the execution of apoptosis. We observed that, compared to controls, Casp3 was up-regulated in the middle region of the mutant gonad, particularly in PGC nuclei (Fig. 3J, K). Altogether, our observations demonstrate that homozygous *pnt^aga^* germ cells die by apoptosis during larval and pupal development, resulting in the final, highly penetrant agametic phenotype. To rule out the possibility that apoptosis is also upregulated in other somatic cells, including the ICs and that PGCs die as a consequence of this, we studied in detail higher magnification views of mutant gonads and found that Caspase3 signal is generally restricted to PGCs (controls: 2.33 ± 0.56 (*n* = 6) apoptotic PGCs and 6.7 ± 1.17 (*n* = 6) apoptotic somatic cells; *pnt^aga^*: 15.13 ± 1.76 (*n* = 8) apoptotic PGCs and 5.75 ± 1.5 (*n* = 8) apoptotic somatic cells; Fig. 3L). From the above results, we conclude that the initial steps of ovary morphogenesis take place correctly and that, as development progresses, *pnt^aga^* germ cells are lost from the mutant gonads. Hence, *pnt^aga^* uncovers a specific function for *pointed* in gonad development.

### *pnt^aga^* affects *pointed* transcription

Using deficiency mapping, genetic tests and genomic sequencing, we mapped the molecular lesion responsible for the *pnt^aga^* phenotype to a two-nucleotide change (AC to CT; FlyBase coordinates: 23,295,576-7) in an intronic region upstream of the TSS of *pnt^short^* (Fig. 4A; Figs. S5, S6). Knowing that the molecular lesion of *pnt^aga^* does not alter the coding sequence of any of the three Pnt proteins, it is likely that *pnt^aga^* causes PGC degeneration by altering transcript levels of the *pnt* locus in the somatic cells of the female gonad. To test this, we performed ddPCR to assess the relative mRNA levels of the different *pnt* isoforms in mutant LL3 larval gonads, and compared them to controls. We observed a significant reduction in the expression levels of both the long and short isoforms *pnt^aga^* (controls: long isoform 22.29 ± 10.8 copies/ng of cDNA; short 11.46 ± 7.5. *pnt^aga^*: long 8.70 ± 0.6; short 3.60 ± 2.1. Two biological and two technical replicates per genotype; Fig. 4B), demonstrating that *pnt^aga^* represents a partial loss-of-function mutation. It impacts *pnt* transcription, leading to altered transcript levels of the different isoforms. Similarly, *pnt*^aga^ mutant embryos showed a decrease in long, intermediate and short isoforms too (controls: long isoform 27.72 ± 2.7 copies/ng of cDNA; intermediate 13.89 ± 1.4; short 28.17 ± 8.5. *pnt^aga^*: long 13.84 ± 3.7; intermediate 5.71 ± 0.8; short 13.47 ± 3.55. Two biological and two technical replicates per genotype; Fig. S4C).

**Figure 4:**
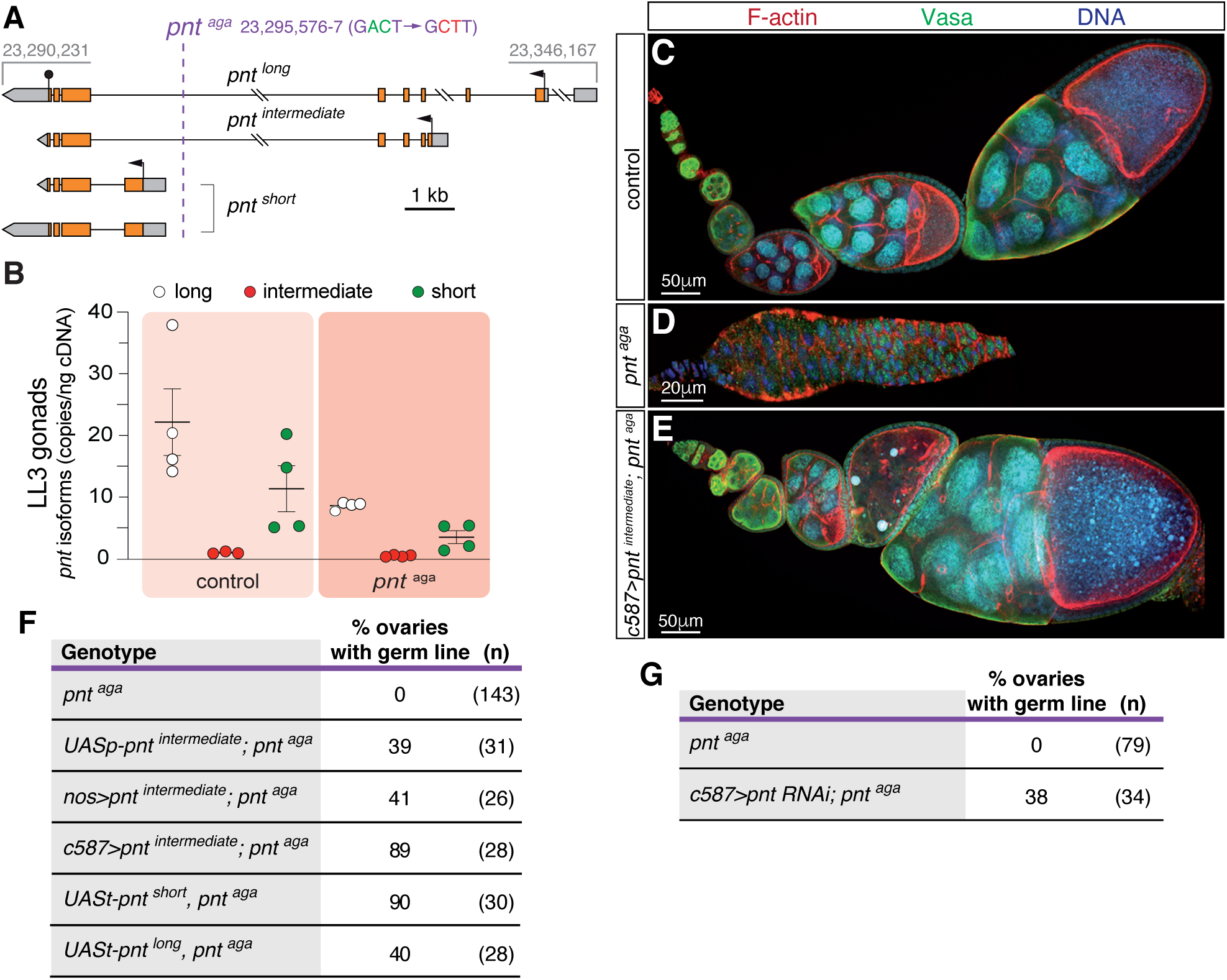
*pnt^aga^* affects *pointed* locus transcription. **(A)** Mapping of the *pnt^aga^* mutation. Numbers refer to genomic coordinates according to FlyBase (release FB2024_02). **(B)** Quantification of the long, intermediate and short *pnt* mRNA levels in control and *pnt^aga^* gonads using droplet-digital PCR. Measurements correspond to three biological replicates and to one or two technical replicates. **(C)** Control, **(D)** *pnt^aga^* and **(E)** *c587>pnt^intermediate^; pnt^aga^* ovarioles stained to visualise F-actin (red), the germ line (anti-Vasa; green) and with a DNA dye (blue). **(F)** Quantification of germline-containing ovaries in different genotypes. Ectopic expression in the soma of any of the *pnt* isoforms rescues the agametic phenotype. **(G)** Quantification of germline-containing ovaries in *pnt^aga^* and in *c587>pnt RNAi; pnt^aga^* females. Reduction of *pnt* function in a *pnt^aga^* background rescues partially the agametic phenotype. Scale bars: 20 or 50 μm, as indicated. Related to Figures S4 and S5.

To determine whether an increase in *pnt* levels could rescue the *pnt^aga^* phenotype, we generated flies containing *UAS-pnt* constructs in a *pnt^aga^* background. We tested the three isoforms, long and short (*UASt-pnt^long^*, *UASt-pnt^short^*; (Klaes et al., 1994)) and intermediate (*UASp-pnt^intermediate^*; this work). In all instances, flies carrying any of the constructs were viable and fertile. Interestingly, *pnt*^aga^ females carrying two copies of any of the *UAS-pnt* constructs showed a significant proportion of ovaries containing germ line (Fig. 4C-F). Since *UAS* constructs can display some basal activity in absence of a *Gal4* driver (Markstein et al., 2008), these phenotypic rescues confirmed the loss-of-function nature of the *pnt^aga^* allele. Next, we determined that *pnt* was required in somatic cells of the gonad, as the somatic *c587-Gal4* driver, but not the germline-specific *nanos-Gal4*, was able to rescue the agametic phenotype when overexpressing *UASp-pnt^intermediate^* (Fig. 4F).

As stated above, the *pnt^aga^* phenotype is opposite to that of *pnt*^RNAi^, the former giving rise to gonads and germaria devoid of germ cells and the latter causing excess germ cell proliferation. Considering that *pnt^aga^* LL3 gonads express reduced levels of long and short *pnt* and that *pnt*^RNAi^ knockdown reduces transcript levels of all isoforms (Fig. S2A), it is possible that germline progression depends on the levels and/or combination of the different *pnt* isoforms expressed. In our working model, a strong reduction in *pnt* levels in somatic cells of the gonad (*c587>pnt*^RNAi^) impairs EGFR signaling and thus results in germ cell overproliferation, while a reduced dose of long and short *pnt* induces germ line disappearance by cell death (*pnt^aga^*). If this hypothesis were right, the *pnt^aga^* phenotype should be suppressed by the removal of all *pnt* function. To test this, we generated flies that carried the *pnt*^RNAi^ construct in a *pnt^aga^* background (*c587>pnt*^RNAi^ + *pnt^aga^*) and scored the germline phenotype in adult females. We found that the expression of *pnt*^RNAi^ rescued partially the agametic phenotype typical of *pnt^aga^*, as experimental *c587>pnt*^RNAi^ + *pnt^aga^* ovaries contained developing egg chambers in 38% of the cases (*n* = 34), whereas control *pnt*^aga^ showed complete absence of germ line (*n* = 79; Fig. 4G). From these results we interpret that germline development relies on the presence of the proper levels of Pnt isoforms — mainly the long and short ones — in the somatic cells of the larval gonad. Thus, the RNAi-induced decrease in *pnt*-mediated signaling causes excess germ cell proliferation, most likely since this would mimic the abrogation of EGFR pathway activity in intermingled cells in the gonad (Gilboa & Lehmann, 2006). Conversely, a milder reduction in the expression of long and short *pnt,* as seen in the *pnt^aga^* condition, induces the opposite effect, germline apoptosis.

### A transcriptomic analysis of *pnt^aga^* gonads identifies cell-type specific changes in cell adhesion, ion binding and membrane activity

Considering the effect of *pnt^aga^* on the various *pointed* isoforms, we aimed to determine how the altered levels of *pointed* transcripts influenced the global transcriptomics of mutant gonads. Using Affymetrix microarrays (GeneChip Drosophila Genome 2.0), we performed a differential gene expression (DGE) analysis of control *vs* mutant LL3 gonads. We identified 253 genes whose expression was changed in *pnt^aga^* gonads, 118 were upregulated and 135 down-regulated (Fig. 5A and Suppl. Data 1; fold change<0.5 or >2; false discovery rate<5%). Gene Ontology (GO) enrichment analyses of the differentially expressed genes yielded a number of terms associated to cell adhesion or ion binding (up-regulated genes) or to membrane activity, including membrane transport, and response to mechanical stimulus (down-regulated genes), among others (Fig. 5B).

**Figure 5:**
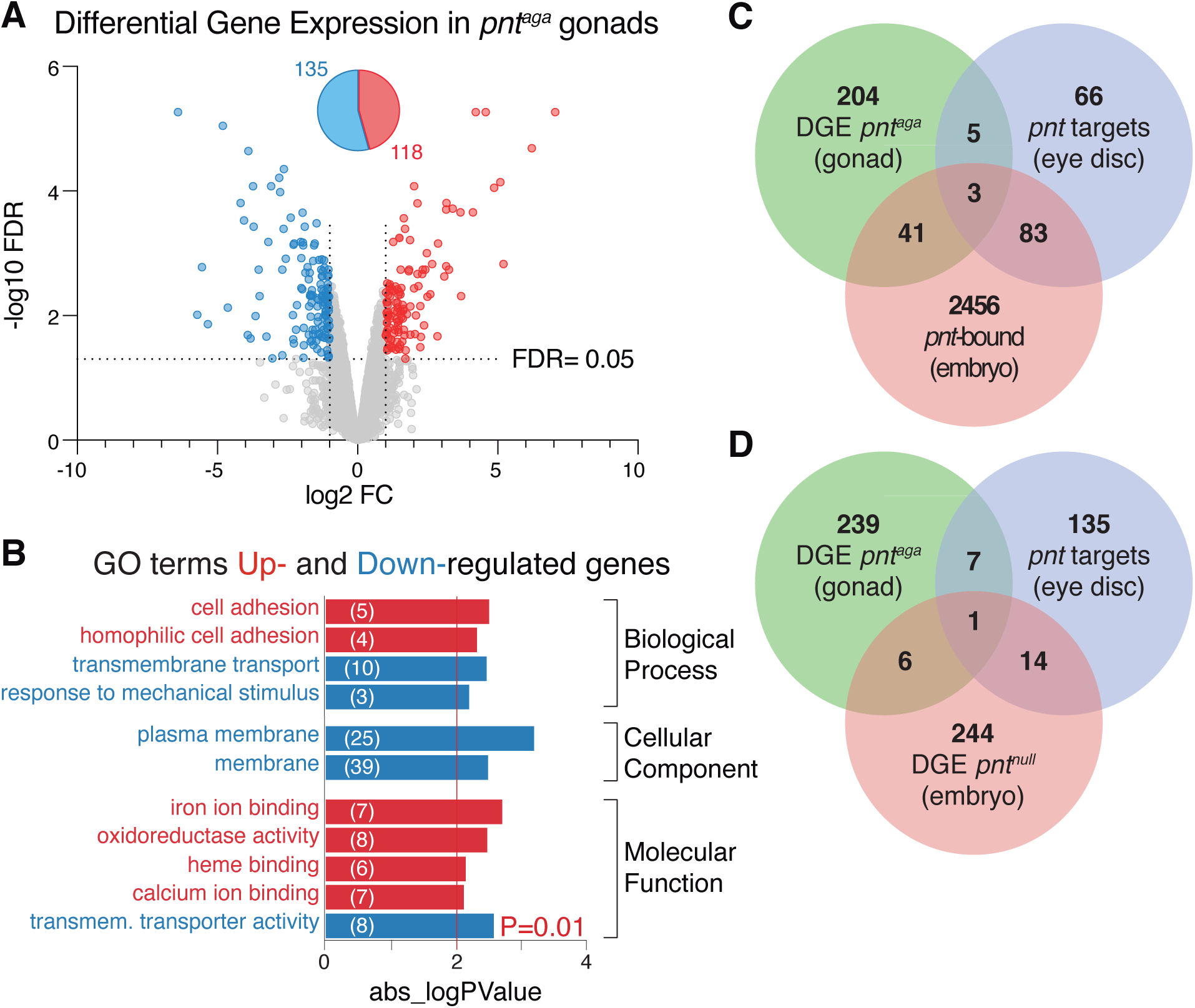
Differential GE in *pnt^aga^* gonads. **(A)** Changes in gene expression (GE) — represented as log2 fold change of *pnt^aga^/*control *versus* the -log10 false discovery rate (FDR) — identified 118 genes upregulated and 135 genes downregulated in *pnt^aga^* compared to controls (linear fold change <0.5 or >2; FDR <5%). **(B)** GO terms identified (p-value<0.01) in the analysis of either up- or down-regulated genes. Numbers in parenthesis correspond to the genes identified in each term. **(C)** Diagram to indicate the number of genes showing differential GE in *pnt^aga^* gonads common to the list of *pnt* targets in the eye disc (Bollepogu Raja et al., 2024) and the genes bound by Pnt in embryos (Webber et al., 2018). **(D)** Diagram representing the number of differentially expressed genes in *pnt^aga^* gonads and in *pnt^null^* embryos, and of direct targets identified in the eye disc. Data related to *pnt*^aga^ correspond to three biological replicates per genotype. Related to Suppl. data S1.

The ETS family of transcriptions factors, comprising Pnt, are versatile in their functions and interact with other transcription factors and cofactors to allow combinatorial control of gene expression. As previously stated, the output of some Ras/MAPK signalling pathways depends upon the biochemical equilibrium between the partners Pnt and Yan, switching from an activation state of high Pnt to a contrasting inactivation state of high Yan. It is important to acknowledge that this bistable model depends on the specific cellular milieu and the intricacies of transcriptional regulation. Hence, a significant proportion of *pnt* targets most likely are tissue specific (Bollepogu Raja et al., 2024; Wu et al., 2020). With this in mind, we compared the list of up- and down-regulated genes in *pnt^aga^* gonads with known *pnt* targets in the eye disc and with *pnt*-bound genes in the embryo (Bollepogu Raja et al., 2024; Webber et al., 2018). We found 8 genes in common with the eye disc targets and 44 also bound by Pnt in the embryo, with only 3 genes common to the three datasets (Fig. 5C; Suppl. Data 1). While the identification of Pnt direct targets in the gonad requires further experimentation, from our results we conclude that the outcome of diminishing *pnt* transcript levels seems to be largely cell-type specific. Finally, considering that the lower levels of long and short *pnt* observed in *pnt^aga^* gonads indicated a partial loss of function situation, we compared the differentially expressed genes in *pnt^aga^* with those identified in *pnt* null mutant embryos (Webber et al., 2018) and identified 7 common genes. Interestingly, 3 of the latter (*hibris, CG1773* and *3-Hydroxymethyl-3-methylglutaryl-CoA lyase*) were up-regulated in *pnt^aga^*, while they were down-regulated in *pnt* null embryos, emphasizing the idea that *pnt*-mediated regulation of gene expression is context-specific. This view is further supported by the limited number (15) of direct *pnt* targets in the eye disc that were also differentially expressed in *pnt* null embryos and by the fact that only 1 gene was common to the three datasets (Fig. 5D; Suppl. Data 1).

## 3. Discussion

Soma-germline interactions are critical to control germline migration, proliferation, the induction and maintenance of germline stem cells, and gamete maturation. A number of pathways regulate gametogenesis, including the EGFR/Ras/MAPK, whose output depends on the activity of the Pnt and Yan cofactors. However, little is known about the role of *pnt* in gametogenesis. Our work, including the isolation and characterization of *pnt*^aga^, demonstrates that *pointed* is required in female somatic gonadal cells to control germline proliferation and survival. For instance, a general reduction in *pnt* mRNAs levels utilising RNA interference results in PGC overproliferation in gonads. On the contrary, the loss-of-function mutation *pnt*^aga^ produces gonads containing fewer PGCs than controls and this reduction in PGC number is due, at least partially, to specific germ cell apoptosis. By the end of L3, mutant female gonads appear almost depleted of PGCs, thus preventing the establishment of germline stem cells. Hence, adult flies become agametic and oogenesis is disrupted. While at present we do not know the molecular mechanisms behind these opposite phenotypes, our work nevertheless highlights a critical function of *pnt* isoform expression in proper gonad development. *pnt*^aga^ maps to an intronic region just upstream of the TSS of the *pnt^short^* isoforms. Based on its close proximity, we speculate that the molecular lesion in *pnt*^aga^ might be affecting the promoter or an enhancer of *pnt^short^* and thus its transcription. If this were the case, since *pnt*^aga^ also affects the transcription of *pnt^long^* in female gonads and of *pnt^intermediate^* and *pnt^long^* in embryos, our analyses suggest that feedback-loop mechanism(s) regulating *pnt* transcription under normal conditions are disrupted in *pnt^aga^*. While we cannot rule out the possibility that *pnt*^aga^ affects a general enhancer for the locus and hence transcription from the three TSSs, because *pnt^aga^* maps very close to *pnt^short^* TSS and because *pnt^short^* seemingly rescues the agametic phenotype more efficiently that *pnt^intermediate^* or *pnt^long^*, we favour the idea that *pnt*^aga^ affects specifically *pnt^short^*. It is known that the three Pnt proteins can perform different functions, as their distinct expression patterns, interactions and transcriptional activities are necessary for proper photoreceptor fate specification and thus eye patterning during larval stages (Wu et al., 2020). These authors demonstrated that Pnt long and intermediate activate *pnt^short^* transcription. However, since they lacked a specific *pnt^short^* mutant, they could not assess regulatory interactions between Pnt short and *pnt^long^* and *pnt^intermediate^*. Our results in embryos and female gonads suggest that Pnt short might be needed to allow proper transcription from the long and intermediate promoters. Our evidence and that of other groups thus point to a complex autoregulatory network involving the three Pnt isoforms to ensure the correct transcriptional activity of the *pnt* locus (O’Neill et al., 1994; Scholz et al., 1993; Wu et al., 2020). This is particularly important in view of the critical function for *pnt* in germline development. Leveraging the drastically opposite phenotypes of a strong loss of function of the *pnt* locus (overproliferation of PGCs) and that of a ∼50% reduction in *pnt^short^* and *pnt^long^* mRNA levels (agametic gonads), it follows that the correct amounts and/or combination of Pnt isoforms are critical to ensure the growth of functional female gonads. Moreover, while we cannot completely rule out the influence of other signalling pathways on *pnt* activity, the fact that *pnt RNAi* mimics EGFR loss-of-function phenotypes strongly suggests that precise regulation of the EGFR signalling pathway is essential for reproduction, at least in females. Although we have not investigated in detail the case of the male gonad and adult testes, the observation that *pnt*^aga^ testes are agametic indicates that this regulatory mechanism may be conserved in males. A similar scenario in which different outcomes for the same signalling pathway depend on the dynamics of pathway activation has been reported in other tissues. For instance, JAK/STAT signaling in the male GSC niche shows contradictory outcomes, such as heterochromatin loss in somatic niche cells or, conversely, activation of heterochromatin in the GSCs (Shi et al., 2006, 2008). In silico simulations predict that transient ligand stimulation, typical in somatic tissues, leads to a transitory decrease in unphosphorylated STAT levels, whereas continuous ligand stimulation, specific of the GSC niche, increases unphosphorylated STAT levels (Li, 2023). Another context in which the distinct expression of transcription factor (TF) isoforms can have profound consequences for tissue function is in cancer disease, where aberrant TFs’ alternative splicing is linked to tumorigenesis. The altered composition of TFs may control different transcriptional programmes, regulate the same target genes with different efficiency or opposite effect, or compete with physiological TF activity (Belluti et al., 2020; Escobar-Hoyos et al., 2019).

The role of *pnt* in controlling embryonic and eye development in *Drosophila* has been known for decades (Brunner et al., 1994; Klaes et al., 1994; Klambt, 1993; Morimoto et al., 1996; Scholz et al., 1993). More recently, novel direct *pnt* targets and the embryonic *pnt/yan* regulatory landscape have been reported (Bollepogu Raja et al., 2024; Webber et al., 2018). *pnt*^aga^ homozygous flies do not show obvious, visible phenotypes during development. Thus, other processes in which *pnt* plays a critical role appear to proceed normally. Considering that the agametic mutation also affects *pnt* transcription in embryos, but seemingly without compromising viability, we surmise that the gonad is particularly sensitive to *pnt* mRNAs levels of the long and short isoforms expressed there. Moreover, our results in the gonad also indicate that the cohort of genes responding to *pnt* activity is context-specific. In fact, there is little overlap between the genes that are regulated by *pnt* in the embryo, the eye disc or the *pnt*^aga^ condition (Bollepogu Raja et al., 2024; Webber et al., 2018). Our study thus identifies a critical function for *pnt* in gonadogenesis and contributes to the description of the *pnt* network in the developing gonad. Importantly, the essential role of *pnt* in gametogenesis seems conserved in mammals, as it parallels known functions of ETS-related proteins in the mammalian testes and ovaries(Chen et al., 2005; Hess et al., 2006; Morrow et al., 2007). In mammals, expression of the ETS related molecule (ERM) is restricted to somatic Sertoli cells and mice with a targeted mutation in ERM fail to maintain spermatogonial stem cells. Self-renewal of the niche seems to be disrupted during the first wave of spermatogenesis with a progressive depletion of the niche, producing a Sertoli-cell-only syndrome (Chen et al., 2005).

While single-cell techniques and bulk RNA sequencing have facilitated the transcriptional characterization of traditionally challenging tissues like the developing larval ovary (Slaidina et al., 2020), these omic approaches have their limitations. For instance, the recently published transcriptomic atlas of various cell types in the developing gonad did not report *pnt* expression in any somatic cells forming the niche or *egfr* transcription in ICs, despite the requirement of EGFR signalling in this cell type for PGC proliferation (Gilboa & Lehmann, 2006; Slaidina et al., 2020). Therefore, direct genetic approaches serve as a valuable complement to broader strategies, aiding in the precise definition of the molecular functions of genes.

## 4. Material and Methods

### Fly stocks and RNAi protocol

We used FlyBase (release FB2024_02) to find information on genes and their functions (Öztürk-Çolak et al., 2024). Flies were grown at 25°C on standard medium unless otherwise stated. The lines used include:

*TM3, twist-Gal4, UAS-nlsGFP* (Halfon et al., 2002)

*UASp-tau::mGFP6* (Bolívar et al., 2006)

*pnt^aga^* (this work)

*pnt^/188^* (Scholz et al., 1993) (Bloomington *Drosophila* Stock Centre (BDSC) #861) P{w[+mC]=lacW} *pnt^1277^* (Scholz et al., 1993)(BDSC #837)

P{w[+mW.hs]=GawB} *pnt^NP5370^* (Kyoto Stock Centre #104978)

*tj-Gal4* (Hayashi et al., 2002)

*c587-Gal4* (Kai & Spradling, 2004)

*nos-Gal4* (Doren et al., 1998) (BDSC #4442)

*UASp-pnt^intermediate^* (this work)

*UASt-pnt.P1 (pnt^short^)* (Klaes et al., 1994) (BDSC #869) *UASt-pnt.P2 (pnt^long^)* (Klaes et al., 1994) (BDSC #399) FRT82B *ubi-nls::GFP* (BDSC #32655)

Df(3R)Exel6280, P{w[+mC]=XP-U}Exel6280 (BDSC #7746) Df(3R)Exel9012, P{w[+mC]=XP-U}Exel9012 (BDSC #7990)

Df(3R)ED6105, P{w[+mW.Scer\FRT.hs3]=3’.RS5+3.3’}ED6105 (Kyoto Stock Center #150336) P{TRiP.JF02227}attp2 (*UAS-pnt RNAi^JF02227^*) (BDSC #31936)

*UAS-pnt RNAi^KK100473^* (Vienna *Drosophila* Resource Center #105390)

The following enhancer-Gal4 constructs from the Janelia Farm collection were used (BDSC stock numbers in parenthesis): *P{GMR44C09-GAL4}attP2* (#41263), *P{GMR43E07-GAL4}attP2* (#45304), *P{GMR44B07-GAL4}attP2* (#45717), *P{GMR46C10-GAL4}attP2* (#46271),

*P{GMR44C01-GAL4}attP2* (#48145), *P{GMR45D11-GAL4}attP2* (#49563), *P{GMR45F08-*

*GAL4}attP2* (#49565), *P{GMR45B10-GAL4}attP2* (#50223), *P{GMR45E10-GAL4}attP2* (#50233).

### *Generation of the* UASp-pnt^intermediate^ *construct*

To express *pnt^intermediate^* in the germline, we first synthesized the *pnt-RD* cDNA from 2-day old control (*y w*) ovaries. mRNA was isolated and purified using the QuickPrep Micro mRNA Purification Kit (GE healthcare) from approx. 100 ovary pairs. 1-2 μg of mRNA were used to synthesized cDNA with 0.5 μg of oligo(dT) (Sigma Genosis) and the Superscript II RNase H Transcriptase (Invitrogen LifeTechnology) in a 20 μl final volume. 2 μl of total cDNA were used in a PCR reaction to amplify the *pnt-RD* cDNA.

The following primers were used:

Forward primer

5’ GCTAGGATCCATGACCAATGAGTGGATCGAT 3’

Reverse primer

5’ GAATGCGGCCGCTTTGCGGTTGGTTGAGTACA 3’

The *pnt-RD* cDNA generated by reverse transcription was then cloned as a BamHI/Not I fragment into pUASp (Rørth, 1998). The resulting construct was verified by restriction digests and sequencing. Transgenic lines were generated by standard procedures.

### Mapping of genomic deficiencies

Genomic DNA from single flies was prepared following the protocol by Georg Dietzl (Barry Dickson’s Laboratory, IMP, Vienna). In short, one fly of the desired genotype was place in a 0.5 ml tube containing 5 µl of squishing buffer and mashed for 5-10 seconds with a pipette tip. After the addition of another 45 µl of squishing buffer, the mix was incubated at room temperature for 30 minutes. The Proteinase K was inactivated by heating the mix at 95°C for 1-2 minutes. Typically, 2 µl of supernatant were used per 20-50 µl PCR reaction.

Squishing buffer: 10 mM Tris-Cl pH 8.2, 1mM EDTA, 25 mM NaCl, 200 µg/ml Proteinase K (enzyme diluted fresh from a frozen stock).

The Exel6280 and Exel9012 deficiencies were generated by FLP/FRT deletions (Parks et al., 2004) and their breakpoints were originally mapped with reasonable precision and reported in FlyBase (flybase.org). They both harbour hybrid elements from the XP5’ (plus and minus) and the WH5’ P-elements, which we used to map the breakpoints in the *pnt* locus accurately.

Internal primers corresponding to the XP elements were paired with genomic primers designed to bind to flaking sequences according to the published breakpoints. The PCR reaction would only amplify a DNA product from the chromosome bearing the deficiency, as internal primers are only complementary to the *P*-element sequences. To sequence the 3’ genomic flanking region of *Df(3R)ED6105* we used a similar strategy, as this deficiency was generated by FLP/FRT-mediated recombination of two RS elements (Ryder et al., 2004). In all cases, genomic DNA was prepared from Deficiency/Balancer flies, diluted 1:25 and used as template in PCR reactions. The primers used were the following (Fig. S6):

*Df(3R)Exel6280*, distal breakpoint:

XP3’+ Flanking genomic region primers:

XP1 forward 5’GCCACCACTTCAAGAACTCTGTAGC 3’ G1 reverse 5’CGAGTTGGCGTTGTTAATGA 3’

*Df(3R) Exel9012*, distal breakpoint:
XP + Flaking genomic region:
52B forward 5’ TTTACTCCAGTCACAGCTTTG 3’
G5 reverse 5’ GCACAGTGGTTGTCATGGTC 3’

*Df(3R) ED6105*, distal breakpoint:
Pry4 (maps to the RS3r 3’ terminal repeat end) forward 5’ CAATCATATCGCTGTCTCACTCA 3’
Genomic reverse 5’ GTTTGTGAGAGGCGGCAGCCA 3’

### *Mapping of* pnt^aga^

Genomic DNA from control *y w* and *pnt^aga^* homozygous flies was prepared as described, diluted 1:25 and used as template in PCR reactions. Genomic primers were designed based on the reference sequence in flybase.org. The following primers were used (Fig. S6):

Forward 5’ AGAACGCGTTGTATGAGGGACCCA 3’

Reverse 5’ GTTTGTGAGAGGCGGCAGCCA 3’

### Sample preparation, Microarray hybridization and data analysis

RNA from 2 larval gonads (approximately 2000-3000 cells) per sample (*y w* controls and *pnt^aga^* homozygous; three biological replicas per genotype) was isolated using magnetic beads. cDNA synthesis, library preparation and amplification (Pico Profiling) are described elsewhere (Gonzalez-Roca et al., 2010). In short, after reverse-transcription each cDNA sample was added to an amplification mix that was subdivided into five equivalent parts for PCR amplification (26 cycles). Amplified cDNA collections were purified with the PureLink PCR Purification Kit (ThermoFisher Scientific), resuspended in 40μl and their concentrations measured using a Nanodrop 1000 spectrophotometer. The six cDNA collections were hybridized to GeneChip *Drosophila* Genome 2.0 Arrays from Affymetrix. Raw data generated with Affymetrix’s AGCC software were used as input for the DTT software. For the resulting datasets, RMA (Robust Microarray Average) normalization and linear modelling by *limma* (Linear Models for Microarray Analysis) was performed. The p-value for the False Discovery Rate (FDR) was ≤ 0.05 and fold change was set at <0.5 or >2.

Homozygous *pnt^aga^* larvae were selected based on their loss of the Tubby marker on the TM6B balancer.

### Detection and Quantification of mRNA levels by ddPCR

We used droplet digital PCR (ddPCR) to quantify mRNA levels of *pnt* long, intermediate and short isoforms from embryos, LL3 gonads and ovary tips (obtained after severing and collecting the germaria and few egg chambers from the whole ovaries). For embryos, RNA was isolated from ∼30 embryos per genotype (*y w* controls and *pnt^aga^* homozygous; three biological replicas each; homozygous *pnt^aga^* embryos were selected based on their loss of the *TM3, twist-Gal4, UAS-GFP* chromosome) with RNAeasy Micro (Qiagen 74004) and QIAshredder (Qiagen 79654) columns. To synthesize cDNA, 10 μl of mRNA were incubated for 5 minutes at 65°C with 1 μl (0.5 μg) of Anchored Oligo(dT)_23_ (Sigma O4387) + 1 μl of 10mM dNTP mix and then placed on ice for 1 minute. 4 μl of 5x First Strand buffer (Y02321) + 1 μl RNaseOUT Recombinant Ribonuclease Inhibitor (40 units/ μl) (100000840) + 2 μl 0.1 M DTT (Y00147) from Invitrogen were added and incubated for 1 minute at 42°C. Next, 1 μl of Super Script II RT from Invitrogen (18064-014) was added and incubated for 50 minutes at 42°C, 15 minutes at 70°C and then placed on ice for 1 minute. Finally, the mix was incubated for 20 minutes at 37°C with 1 μl of RNase H from Invitrogen (18021-014). For LL3 gonads, see above. Finally, to determine *pnt* expression in ovary tips, we dissected 30 adult ovaries per genotype and manually severed off the tips (which included the germarium and few young egg chambers per ovariole) using tungsten needles. RNA was isolated as in the case of embryos.

Primers used were the following (5’-3’): *pnt-RB* (long) isoform, F (exon 2): TTCTGTCCAGCCTAGTTGAG, R (exon 5): AGTACCTCGTTGACCTTGCG; *pnt-RD* (intermediate) isoform, F (exon 4): AACCACCAATCATAGCAAGC, R (exon 5): AGTACCTCGTTGACCTTGCG; and *pnt-RC/E* (short) isoform, F (exon 8): TGCGAATGCCTACTACACGG, R (exon 9): TACCGCTGCCATTGACAGTC. Ribosomal protein L32 (*RpL32*) and β-Tubulin (β*-Tub*) were used for normalization. *RpL32*, F: ATGACCATCCGCCCAGCATAC, R: GCTTAGCATATCGATCCGACTGG; β*-Tub*, F: GCAGTTCACCGCTATGTTCA , R: CGGACACCAGATCGTTCAT.

For the ddPCR, each reaction contained 10 μl of Master Mix ddPCR EVAGreen (Bio-Rad), 150 nM of each primer and 2,5 ng of cDNA template (except in the case of the housekeeping control, where we used 0,5 ng). Samples were prepared in duplicate with 10% additional volumen. Droplet generation, PCR amplification and droplet analysis were done using the QX200 AutoDG ddPCR system (Bio-Rad). PCR conditions were: 95°C for 5 minutes (1x), 95°C for 1 minute and 65.2°C for 2 minutes (44x). For all steps a ramp rate of 2°C/s was used. Data were analysed with the Quantasoft software 1.7.4.0917 (Bio-Rad).

### Developmental staging of larvae and pupae

At 25°C, the end of embryogenesis is 24 hours after egg laying (AEL). The first and second larval instars are 24 hours each. Third larval instar lasts for 48 hours and puparium formation starts 120 hours AEL. In this study, ML3 (mid-third instar larvae) refers to larvae that crawled out of the food but that have not initiated pupariation. At this stage, most of TFs are still forming and cap cells are not ready visible. LL3 (late-third instar larvae) is used to identify larvae about to initiate pupariation. Their gonads present a number of already-formed TFs and some cap cells can be recognised. The white pupa stage marks the initiation of pupariation and it is characterised by pale and clear pupae. At this stage, the GSC niches with their respective TFs and cap cell rosettes are well established (Yatsenko & Shcherbata, 2018).

### Immunohistochemistry

Adult flies were yeasted for 2 days before dissection in cold Ringer’s. Ovaries and testes were then fixed for 10 minutes in 6% formaldehyde in fix buffer (600μl of heptane were added per 100μl of fix). After fixing, samples were permeabilized in 1% Triton in PBS for 2 hours and then blocked for 1 hour with PAT before overnight incubation with primary antibodies in PAT. The following morning primary antibodies were rinsed with PAT and the samples washed three times with PBT for 30 minutes. The incubation with secondary antibodies was done in PBT during 2 to 4 hours. Ovaries and testes were then washed three times with PTW, 10 minutes each. Mounting of testes and individual ovarioles was done in Vectashield medium (Vector Laboratories; Cat# H1000; RRID: AB_2336789).

To stain larval gonads, gonads embedded in the fat body were fixed in 5% formaldehyde in Ringer’s for 20 minutes then washed for 5, 10 and 45 minutes with 1% PBT. The fat body was then blocked in 0.3% PBTB for 1 hour with gentle agitation. After blocking, the tissue was incubated overnight with the desired primary antibody diluted in 0.3% PBTB at 4°C with gentle agitation. The next day the fat body was washed thrice (30 minutes each) with 0.3% PBTB. After blocking with 0.3% PBTB supplemented with 5% fetal bovine serum for 1 hour, the fat body was incubated for 2 hours with the secondary antibodies in blocking solution, followed by three washes of 30 min each with 0.3% PBT. Gonads were dissected from the fat body and mounted in Vectashield.

Embryos collected overnight on yeasted agar plates were dechorionated with bleach for 2 minutes and fixed for 20 minutes in 2 ml of 4% formaldehyde in PBS with an equal volume of heptane. After fixing, the aqueous phase was removed and 2 ml of methanol were added to remove the vitelline membrane. Devitellinized embryos were rinsed in methanol thrice and stored at -20°C. For antibody staining, embryos were re-hydrated in PBS:methanol for 5 minutes and then washed in PBS, in PBS:PAT and finally rinsed in PAT twice. Next, they were blocked for 1-2 hours in PAT at 4°C. Primary antibody incubation was done overnight at 4°C in PAT, followed by a blocking step for several hours in PBT supplemented with 4% goat serum prior to secondary antibody incubation for 2-4 hours in PBT. Finally, embryos were washed thrice with PBT, 10 minutes each, and mounted in Vectashield.

Primary antibodies were used at the following concentrations: mouse anti-Hts (1B1, Developmental Studies Hybridoma Bank (DSHB) Cat# 1b1, RRID:AB_528070), 1:100; rabbit anti-Vasa 1:2000 (a gift from R. Lehmann); rat anti-*D*E-Cadherin (DSHB Cat# DCAD2, RRID:AB_528120) 1:100; mouse monoclonal anti-Engrailed (DSHB Cat# 4D9; RRID: AB_528224) 1:10; rat anti-Bab2 (a gift from F. Laski) 1:4000; rabbit anti-cleaved-Caspase3 (from Cell Signalling Technology) 1:50; goat anti-GFP FITC-conjugated (Abcam Cat# ab6662, RRID:AB_305635) 1:500. Secondary antibodies Alexa-Fluor 488, Cy2, Cy3 and Cy5 (Jackson Immuno Research Laboratories, Inc.; final concentrations of 1:100) and conjugated anti-GFP-488 nanobody (alpaca anti-GFP-Booster_Atto488 from ChromoTek, Cat# gba488-100, RRID:AB_2631386; final concentration 1:200) were incubated for four hours. To stain DNA, ovaries were incubated for 10 minutes with Hoechst (Sigma B2883 10 mg/ml in H_2_O). To visualise F-actin, fixed ovaries were incubated in PBT + Rhodamine-labelled Phalloidin (Biotium Cat# BT-00027; final concentration 1:200) for 20 minutes.

Ringer’s: 128 mM NaCl, 2 mM KCl, 1.8mM CaCl2, 4 mM MgCl2, 35.5 mM Sucrose, 5 mM Hepes pH 6.9

Fix buffer: 16.7 mM KPO4 pH 6.8, 75 mM KCl, 25 mM NaCl and 3.3 mM MgCl2

PAT: PBS, 1% BSA, 0.1% Triton, 0.05% azide

PBT: PBS, 0.1 BSA, 0.1% tween20 PTW: PBS, 0.1% tween20

1% PBT: 1% Triton x-100 in PBS

0.3% PBTB: 0.3% Triton X-100 and 1% BSA in PBS

### Imaging, processing and quantification of larval, pupal and adult samples

Images were acquired with Leica’s TCS-SP5 or Stellaris confocal microscopes, analysed utilising ImageJ, and processed with Adobe Photoshop and Adobe Illustrator. Z stacks of fixed samples were taken at 0.7 μm intervals using 40x/1.3 and 63x/1.4 NA oil immersion objectives.

Quantification of the number of Vasa+ cells was done as described in (Rosales-Nieves et al., 2023). Quantification of the number of Casp3+ cells was done manually using the *multi-point* tool in ImageJ.

### Statistical Analysis

Experiments involving gonads or ovaries were performed with at least three biological replicates. Samples were collected from at least 5 different larvae, white pupae or adult females grown in equivalent environmental conditions. For each of the quantifications, the arithmetic mean and the standard error of the mean (SEM) of the different experimental settings are shown. Sample sizes correspond to the number of gonads or adult ovaries analysed. Statistically significant differences between control and experimental samples were calculated with a Student’s *t*-test. *= p<0.05; **=p<0.005; ***=p<0.0005; ****=p<0.0001. Only significant differences are indicated in the graphs.

The quantifications of mRNA concentrations using digital PCR were performed with a minimum of two biological replicates.

### Experimental genotypes

<u>Figure 1</u>

(F, G) pnt enhancer 45E10>tau::GFP: y w/w; P{GMR45E10-GAL4}attP2/+; UASp-tau::mGFP6/+

<u>Figure 2</u>

(A, D) control: w, c587-Gal4/y w;; UASp-tau::mGFP6/+

(B, E) *c587>pnt RNAi: w, c587-Gal4/y v;;* P{TRiP.JF02227}attp2*/+*

(C) control: w, c587-Gal4/w; +/CyO

*c587>pnt RNAi (JF02227): w, c587-Gal4/y v;;* P{TRiP.JF02227}attp2*/+ c587>pnt RNAi (KK100473): w, c587-Gal4/w; UAS-pnt RNAi^KK100473^/+*

<u>Figure 3</u>

(A-L) control: w;; FRT-82B cu sr pnt^aga^/TM3, twist-Gal4, UAS-nlsGFP pnt^aga^: w;; FRT-82B cu sr pnt^aga^

<u>Figure 4</u>

(B-G) control: w;; FRT-82B cu sr pnt^aga^/TM3, twist-Gal4, UAS-nlsGFP pnt^aga^: w;; FRT-82B cu sr pnt^aga^

(E, F) *UASp-pnt^intermediate^; pnt^aga^: UASp-pnt^intermediate^; FRT-82B cu sr pnt^aga^*

*nos>pnt^intermediate^; pnt^aga^: y w/w; nos-Gal4/UASp-pnt^intermediate^; FRT-82B cu sr pnt^aga^ c587>pnt^intermediate^; pnt^aga^: w, c587-Gal4/w; UASp-pnt^intermediate^; FRT-82B cu sr pnt^aga^ UASp-pnt^intermediate^; pnt^aga^: UASp-pnt^intermediate^; FRT-82B cu sr pnt^aga^*

*UASp-pnt^short^ pnt^aga^: UASp-pnt^short^, sr pnt^aga^ UASp-pnt^long^ pnt^aga^: UASp-pnt^long^, sr pnt^aga^*

*c587>pnt RNAi; pnt^aga^: w, c587-Gal4/w; UAS-pnt RNAi^KK100473^/+; FRT-82B cu sr pnt^aga^*

<u>Figure 5</u>

control: y *w*

*pnt^aga^: w;; FRT-82B cu sr pnt^aga^*

<u>Figure S1</u>

(A-C) *NP5370>tau::GFP: y w;;* P{w[+mW.hs]=GawB} *pnt^NP5370^*/*UASp-tau::mGFP6*

(D-L) *pnt* enhancer*>tau::GFP: y w/w; P{GMR* enhancer*-GAL4}attP2/+; UASp-tau::mGFP6/+*

The nine enhancers used are listed in the *Fly stocks* section above.

<u>Figure S2</u>

(B) control*: w; +/CyO*

*tj>pnt RNAi (JF02227): w/y v; tj-Gal4/+;* P{TRiP.JF02227}attp2*/+ tj>pnt RNAi (KK100473): w; UAS-pnt RNAi^KK100473^/tj-Gal4*

(C) tj>tau::GFP: w/y w; tj-Gal4/+; UASp-tau::mGFP6/+

(D, E) *c587>tau::GFP: w, c587-Gal4/y w;; UASp-tau::mGFP6/+*

(F, G) *nos>tau::GFP: w/y w; nos-Gal4/+; UASp-tau::mGFP6/+*

<u>Figures S3 and S4</u>

(A-C) control: y *w*

*pnt^aga^: w;; FRT-82B cu sr pnt^aga^*

<u>Figure S5</u>

(B, C) control: *w*

*pnt^aga^: w;; FRT-82B cu sr pnt^aga^*

(D) *pnt^/188^*/Df(3R)6280: *y w f/w;; FRT-82B pnt^/188^/*Df(3R)Exel6280, P{w[+mC]=XP-U}Exel6280 *pnt^/188^*/Df(3R)9012: *y w f/w;; FRT-82B pnt^/188^/*Df(3R)Exel9012, P{w[+mC]=XP-U}Exel9012 *pnt^/188^*/Df(3R)ED6105: *y w f/w;; FRT-82B pnt^/188^/*Df(3R)ED6105,

P{w[+mW.Scer\FRT.hs3]=3’.RS5+3.3’}ED6105

*pnt^aga^*/ *pnt^/188^*: *w/y w f;; FRT-82B cu sr pnt^aga^*/*FRT-82B pnt^/188^ pnt^aga^*/*pnt^1277^*: *w;; FRT-82B cu sr pnt^aga^*/ P{w[+mC]=lacW} *pnt^1277^*

*pnt^aga^*/Df(3R)9012: *w;; FRT-82B cu sr pnt^aga^*/Df(3R)Exel9012, P{w[+mC]=XP-U}Exel9012 *pnt^aga^*/Df(3R)6280: *w;; FRT-82B cu sr pnt^aga^*/Df(3R)Exel6280, P{w[+mC]=XP-U}Exel6280 *pnt^aga^*/Df(3R)ED6105: *w;; FRT-82B cu sr pnt^aga^*/Df(3R)ED6105, P{w[+mW.Scer\FRT.hs3]=3’.RS5+3.3’}ED6105

## Acknowledgments

We thank the BDSC, the VDRC and the DSHB (University of Iowa) for DNAs, fly stocks and antibodies. The help of Susan Parkhurst and members of her group for discussions and technical support is also acknowledged. We thank members of our group, M.D. Martín-Bermudo, C. Huertas, J. J. Pérez-Moreno, S.C. Herrera, and J. Hombría for comments on the manuscript and helpful discussions. We are grateful to the scientific-technical facilities at CABD for expert support.

## Author Contributions

AERN and AGR conceived and designed research; AERN, MMM and LLO performed research; AERN, JGM and AGR analysed data and wrote the paper.

## Figure Legends

**Figure S1: Expression patterns of several regulatory elements in the *pointed* locus of LL3 gonads and adult ovaries.** In all cases, the regulatory elements drive expression of the Gal4 transcriptional activator. As a reporter, we used the *UASp-tau::GFP* construct. **(A-C)** LL3 gonad, germarium and stage 6 egg chamber of *NP5370>tau::GFP* female larva and adult. They have been stained with anti-Hts (red; to visualise cell outlines and germline spectrosomes and fusomes), anti-Vasa (green; to label the germ line), anti-GFP (white; to show Tau::GFP localization) and with a DNA dye (blue; to mark chromatin). The molecular mapping of the NP5370 insertion is represented in the genomic map above the set of panels. **(D-L)** LL3 gonads and germaria of nine different *pnt* enhancer*>tau::GFP* female larvae and adults. They have been stained with anti-Hts (red), anti-Vasa (green) and anti-GFP (white). GSC: germline stem cell; TFC: terminal filament cell; CpC: cap cell; IC: intermingled somatic cell; EC: escort cell; ShC: sheath cell; SwC: Swarm cell. Scale bars: 20 μm. Related to Figure 1.

**Figure S2: *pointed* RNAi lines and *Gal4* drivers used in this study. (A)** Representation of the exon-intron organization of the *pnt* locus. The two RNAi lines used in this work target similar fragments of the coding region common to all of the transcripts. Numbers of genomic coordinates according to FlyBase (release FB2024_02). **(B)** Quantification of the number of Vasa+ cells in control, *tj>pntRNAi^JF02227^* and *tj>pntRNAi^KK100473^* LL3 gonads. The arithmetic mean and the SEM are shown for each of the genotypes. **(C)** Pattern of expression of the *tj-Gal4* line in control gonads visualised by the distribution of the Tau::GFP reporter (white). **(D, E)** Pattern of expression of the *c587-Gal4* line in control LL3 gonads and adult germaria as shown by the Tau::GFP reporter. The localization of the Hts (red) and Vasa (green) proteins is used to outline cell shapes and to label the germ line, respectively. **(F, G)** Pattern of expression of the *nos-Gal4* line in control LL3 gonads and adult germaria as shown by the Tau::GFP reporter. The localization of Hts (red) is used to outline cell shapes. *=p<0.05; **=p<0.005. Scale bar: 20 μm. Related to Figure 2.

**Figure S3: *pnt^aga^* gives rise to agametic testes. (A, B)** dark field images of control (A) and *pnt^aga^* (B) testes. **(A’, B’)** Control (A’) and *pnt^aga^* (B’) testes stained to visualize F-actin (red) and Vasa (green; to label the germ line). *pnt^aga^* testes are devoid of germline cells. Scale bars: 20 μm. Related to Figure 3.

**Figure S4: *pnt^aga^* does not affect PGC numbers in embryonic gonads. (A, B)** Control (A) and *pnt^aga^* (B) embryos at different stages of embryogenesis stained with *D*E-cad (red; to label cell outlines) and Vasa (green; to mark PGCs). **(C)** Quantification of the long, intermediate and short *pnt* mRNA levels in control and *pnt^aga^* embryos using droplet-digital PCR. Measurements correspond to two biological and two technical replicates. In spite of no obvious differences in PGC numbers in both genotypes, *pnt* mRNA levels are decreased in *pnt^aga^* embryos. Scale bars: 50 μm. Related to Figure 3.

**Figure S5: Molecular mapping of *pnt^aga^*. (A)** Scheme showing the molecular characterization of *pnt^aga^*, *pnt^/188^*, *pnt^1277^* and the three deficiencies used to map *pnt^aga^*. Numbers refer to genomic coordinates according to FlyBase (release FB2024_02). Mapping of *pnt^1277^* and the *pnt^/188^* deletion according to (Scholz et al., 1993). **(B)** Sequence of the region around the mapped *pnt^aga^* mutation. The “subject” sequence corresponds to the reference sequence in FlyBase (release FB2024_02). The control “query” sequence is that of *y w* flies. **(C)** Chromatogram of the relevant region showing the two-base pair change in *pnt^aga^*. **(D)** Table summarising the genetic characterization of different combinations of *pointed* mutants and deficiencies.

**Figure S6: Primers used to map *pnt^aga^* and associated deficiencies.** The precise mapping of Df(3R)Exel9012, Df(3R)Exel6280 and Df(3R)ED6105 were aided by the transposons used to generate them in the first place. Numbers in parenthesis correspond to the genomic coordinates of the breakpoints. Also shown are the genomic position of the primers used to map *pnt^aga^* and the genomic coordinates of the two-base pair substitution found in this mutant.

## References

Belluti, S., Rigillo, G., & Imbriano, C. (2020). Transcription factors in cancer: When alternative splicing determines opposite cell fates. In Cells (Vol. 9, Issue 3). Multidisciplinary Digital Publishing Institute (MDPI). 10.3390/cells9030760

Boisclair Lachance, J. F., Peláez, N., Cassidy, J. J., Webber, J. L., Rebay, I., & Carthew, R. W. (2014). A comparative study of pointed and yan expression reveals new complexity to the transcriptional networks downstream of receptor tyrosine kinase signaling. Developmental Biology, 385(2), 263–278. 10.1016/j.ydbio.2013.11.002

Bolívar, J., Pearson, J., López-Onieva, L., & González-Reyes, A. (2006). Genetic dissection of a stem cell niche: The case of the Drosophila ovary. Developmental Dynamics, 235(11), 2969–2979. 10.1002/dvdy.20967

Bollepogu Raja, K. K., Yeung, K., Shim, Y. K., & Mardon, G. (2024). Integrative genomic analyses reveal putative cell type-specific targets of the Drosophila ets transcription factor Pointed. BMC Genomics, 25(1). 10.1186/s12864-024-10017-7

Brunner, D., Dücker, K., Oellers, N., Hafen, E., Scholzi, H., & Klambt, C. (1994). The ETS domain protein Pointed-P2 is a target of MAP kinase in the Sevenless signal transduction pathway. Nature, 370(6488), 386–389. 10.1038/370386a0

Chen, C., Ouyang, W., Grigura, V., Zhou, Q., Carnes, K., Lim, H., Zhao, G. Q., Arber, S., Kurpios, N., Murphy, T. L., Cheng, A. M., Hassell, J. A., Chandrashekar, V., Hofmann, M. C., Hess, R. A., & Murphy, K. M. (2005). ERM is required for transcriptional control of the spermatogonial stem cell niche. Nature, 436(7053), 1030–1034. 10.1038/nature03894

Dittmer, J., & Nordheim, A. (1998). Ets transcription factors and human disease. Biochimica et Biophysica Acta (BBA) - Reviews on Cancer, 1377(2), F1–F11. 10.1016/S0304-419X(97)00039-5

Doren, M. Van, Williamson, A. L., & Lehmann, R. (1998). Regulation of zygotic gene expression in Drosophila primordial germ cells. Current Biology, 8(4), 243–246. 10.1016/S0960-9822(98)70091-0

Escobar-Hoyos, L., Knorr, K., & Abdel-Wahab, O. (2019). Aberrant RNA splicing in cancer. In Annual Review of Cancer Biology (Vol. 3, Issue 1, pp. 167–185). Annual Reviews Inc. 10.1146/annurev-cancerbio-030617-050407

Gilboa, L., & Lehmann, R. (2006). Soma-germline interactions coordinate homeostasis and growth in the Drosophila gonad. Nature, 443(7107), 97–100. 10.1038/nature05068

Godt, D., & Laski, F. A. (1995). Mechanisms of cell rearrangement and cell recruitment in *Drosophila* ovary morphogenesis and the requirement of *bric à brac*. Development, 121(1), 173–187. 10.1242/dev.121.1.173

Gonzalez-Roca, E., Garcia-Albéniz, X., Rodriguez-Mulero, S., Gomis, R. R., Kornacker, K., & Auer, H. (2010). Accurate expression profiling of very small cell populations. PLoS ONE, 5(12). 10.1371/journal.pone.0014418

Halfon, M. S., Gisselbrecht, S., Lu, J., Estrada, B., Keshishian, H., & Michelson, A. M. (2002). New fluorescent protein reporters for use with the Drosophila Gal4 expression system and for vital detection of balancer chromosomes. Genesis (United States), 34(1–2), 135–138. 10.1002/gene.10136

Hayashi, S., Ito, K., Sado, Y., Taniguchi, M., Akimoto, A., Takeuchi, H., Aigaki, T., Matsuzaki, F., Nakagoshi, H., Tanimura, T., Ueda, R., Uemura, T., Yoshihara, M., & Goto, S. (2002). GETDB, a database compiling expression patterns and molecular locations of a collection of Gal4 enhancer traps. Genesis (United States), 34(1–2), 58–61. 10.1002/gene.10137

Hess, R. A., Cooke, P. S., Hofmann, M.-C., & Murphy, K. M. (2006). Mechanistic Insights into the Regulation of the Spermatogonial Stem Cell Niche. Cell Cycle, 5(11), 1164– 1170. 10.4161/cc.5.11.2775

Hollenhorst, P. C., McIntosh, L. P., & Graves, B. J. (2011). Genomic and biochemical insights into the specificity of ETS transcription factors. Annual Review of Biochemistry, 80, 437–471. 10.1146/annurev.biochem.79.081507.103945

Hsu, T., & Schulz, R. A. (2000). Sequence and functional properties of Ets genes in the model organism Drosophila. Oncogene, 19(55), 6409–6416. 10.1038/sj.onc.1204033

Kai, T., & Spradling, A. (2003). An empty *Drosophila* stem cell niche reactivates the proliferation of ectopic cells. Proceedings of the National Academy of Sciences, 100(8), 4633–4638. 10.1073/pnas.0830856100

Kai, T., & Spradling, A. (2004). Differentiating germ cells can revert into functional stem cells in Drosophila melanogaster ovaries. Nature, 428(6982), 564–569. 10.1038/nature02436

Klaes, A., Menne, T., Stollewerk, A., Scholz, H., & Klambt, C. (1994). The Ets Transcription Factors Encoded by the Drosophila Gene pointed Direct Glial Cell Differentiation in the Embryonic CNS. In Cell (Vol. 76).

Klambt, C. (1993). The Drosophila gene pointed encodes two ETS-like proteins which are involved in the development of the midline glial cells. Development, 117(1), 163–176. 10.1242/dev.117.1.163

Lee, T., & Montell, D. J. (1997). Multiple Ras Signals Pattern the Drosophila Ovarian Follicle Cells. In DEVELOPMENTAL BIOLOGY (Vol. 185).

Li, W. X. (2023). Computational simulation of JAK/STAT signaling in somatic versus germline stem cells. Developmental Dynamics. 10.1002/dvdy.684

Liu, M., Lim, T. M., & Cai, Y. (2010). The Drosophila female germline stem cell lineage acts to spatially restrict DPP function within the niche. Science Signaling, 3(132). 10.1126/scisignal.2000740

Margolis, J., & Spradling, A. (1995). Identification and behavior of epithelial stem cells in the *Drosophila* ovary. Development, 121(11), 3797–3807. 10.1242/dev.121.11.3797

Markstein, M., Pitsouli, C., Celniker, S. E., & Perrimon, N. (2008). Exploiting position effects and the gypsy retrovirus insulator to engineer precisely expressed transgenes. http://www.nature.com/naturegenetics

Matsuoka, S., Hiromi, Y., & Asaoka, M. (2013). Egfr signaling controls the size of the stem cell precursor pool in the Drosophila ovary. Mechanisms of Development, 130(4–5), 241–253. 10.1016/j.mod.2013.01.002

Meignin, C., Alvarez-Garcia, I., Davis, I., & Palacios, I. M. (2007). The Salvador-Warts-Hippo Pathway Is Required for Epithelial Proliferation and Axis Specification in Drosophila. Current Biology, 17(21), 1871–1878. 10.1016/j.cub.2007.09.062

Morimoto, A. M., Jordan, K. C., Tietze, K., Britton, J. S., O’Neill, E. M., & Ruohola-Baker, H. (1996). Pointed, an ETS domain transcription factor, negatively regulates the EGF receptor pathway in Drosophila oogenesis. Development, 122(12), 3745–3754. 10.1242/dev.122.12.3745

Morrow, C. M. K., Hostetler, C. E., Griswold, M. D., Hofmann, M. C., Murphy, K. M., Cooke, P. S., & Hess, R. A. (2007). ETV5 is required for continuous spermatogenesis in adult mice and may mediate blood-testes barrier function and testicular immune privilege. Annals of the New York Academy of Sciences, 1120, 144–151. 10.1196/annals.1411.005

Oikawa, T., & Yamada, T. (2003). Molecular biology of the Ets family of transcription factors. Gene, 303, 11–34. 10.1016/S0378-1119(02)01156-3

O’Neill, E. M., Rebay, I., Tjian, R., & Rubin, G. M. (1994). The activities of two Ets-related transcription factors required for drosophila eye development are modulated by the Ras/MAPK pathway. Cell, 78(1), 137–147. 10.1016/0092-8674(94)90580-0

Öztürk-Çolak, A., Marygold, S. J., Antonazzo, G., Attrill, H., Goutte-Gattat, D., Jenkins, V. K., Matthews, B. B., Millburn, G., dos Santos, G., & Tabone, C. J. (2024). FlyBase: updates to the Drosophila genes and genomes database. Genetics, 227(1). 10.1093/genetics/iyad211

Parks, A. L., Cook, K. R., Belvin, M., Dompe, N. A., Fawcett, R., Huppert, K., Tan, L. R., Winter, C. G., Bogart, K. P., Deal, J. E., Deal-Herr, M. E., Grant, D., Marcinko, M., Miyazaki, W. Y., Robertson, S., Shaw, K. J., Tabios, M., Vysotskaia, V., Zhao, L., … Francis-Lang, H. L. (2004). Systematic generation of high-resolution deletion coverage of the Drosophila melanogaster genome. Nature Genetics, 36(3), 288–292. 10.1038/ng1312

Rørth, P. (1998). Gal4 in the Drosophila female germline.

Rosales-Nieves, A. E., Marín-Menguiano, M., Campoy-Lopez, A., & González-Reyes, A. (2023). Visualization and Quantification of Drosophila Larval Ovaries (pp. 37–47). 10.1007/978-1-0716-2970-3_2

Ryder, E., Blows, F., Ashburner, M., Bautista-Llacer, R., Coulson, D., Drummond, J., Webster, J., Gubb, D., Gunton, N., Johnson, G., O’Kane, C. J., Huen, D., Sharma, P., Asztalos, Z., Baisch, H., Schulze, J., Kube, M., Kittlaus, K., Reuter, G., … Russell, S. (2004). The DrosDel collection: A set of P-element insertions for generating custom chromosomal aberrations in Drosophila melanogaster. Genetics, 167(2), 797–813. 10.1534/genetics.104.026658

Sahut-Barnola, I., Godt, D., Laski, F. A., & Couderc, J.-L. (1995). Drosophila Ovary Morphogenesis: Analysis of Terminal Filament Formation and Identification of a Gene Required for This Process. Developmental Biology, 170(1), 127–135. 10.1006/dbio.1995.1201

Scholz, H., Deatrick, J., Klaes, A., & Klambt, C. (1993). Genetic Dissection of pointed, a Drosophila Gene Encoding Two ETS-Related Proteins.

Sharrocks, A. D. (2001). The ETS-domain transcription factor family. Nature Reviews Molecular Cell Biology, 2(11), 827–837. 10.1038/35099076

Shi, S., Calhoun, H. C., Xia, F., Li, J., Le, L., & Li, W. X. (2006). JAK signaling globally counteracts heterochromatic gene silencing. Nature Genetics, 38(9), 1071–1076. 10.1038/ng1860

Shi, S., Larson, K., Guo, D., Lim, S. J., Dutta, P., Yan, S. J., & Li, W. X. (2008). Drosophila STAT is required for directly maintaining HP1 localization and heterochromatin stability. Nature Cell Biology, 10(4), 489–496. 10.1038/ncb1713

Slaidina, M., Banisch, T. U., Gupta, S., & Lehmann, R. (2020). A single-cell atlas of the developing Drosophila ovary identifies follicle stem cell progenitors. Genes and Development, 34(3), 239–249. 10.1101/gad.330464.119

Verger, A., & Duterque-Coquillaud, M. (2002). When Ets transcription factors meet their partners alexis verger and martine duterque-coquillaud. In BioEssays (Vol. 24, Issue 4, pp. 362–370). 10.1002/bies.10068

Vivekanand, P., Tootle, T. L., & Rebay, I. (2004). MAE, a dual regulator of the EGFR signaling pathway, is a target of the Ets transcription factors PNT and YAN. Mechanisms of Development, 121(12), 1469–1479. 10.1016/j.mod.2004.07.009

Wang, Y., Huang, Z., Sun, M., Huang, W., & Xia, L. (2023). ETS transcription factors: Multifaceted players from cancer progression to tumor immunity. In Biochimica et Biophysica Acta - Reviews on Cancer (Vol. 1878, Issue 3). Elsevier B.V. 10.1016/j.bbcan.2023.188872

Webber, J. L., Zhang, J., Massey, A., Sanchez-Luege, N., & Rebay, I. (2018). Collaborative repressive action of the antagonistic ETS transcription factors Pointed and Yan fine-tunes gene expression to confer robustness in Drosophila. Development (Cambridge), 145(13). 10.1242/dev.165985

Wu, C., Lachance, J. F. B., Ludwig, M. Z., & Rebay, I. (2020). A context-dependent bifurcation in the Pointed transcriptional effector network contributes specificity and robustness to retinal cell fate acquisition. PLoS Genetics, 16(11). 10.1371/journal.pgen.1009216

Yatsenko, A. S., & Shcherbata, H. R. (2018). Stereotypical architecture of the stem cell niche is spatiotemporally established by miR-125-dependent coordination of Notch and steroid signaling. Development (Cambridge), 145(3). 10.1242/dev.159178

Zhu, C. H., & Xie, T. (2003). Clonal expansion of ovarian germline stem cells during niche formation in Drosophila. In Development (Vol. 130, Issue 12, pp. 2579–2588). 10.1242/dev.00499

