## Supplementary figures S1-S6 for "ETS TRANSCRIPTION FACTOR POINTED CONTROLS GERMLINE SURVIVAL IN *DROSOPHILA*"

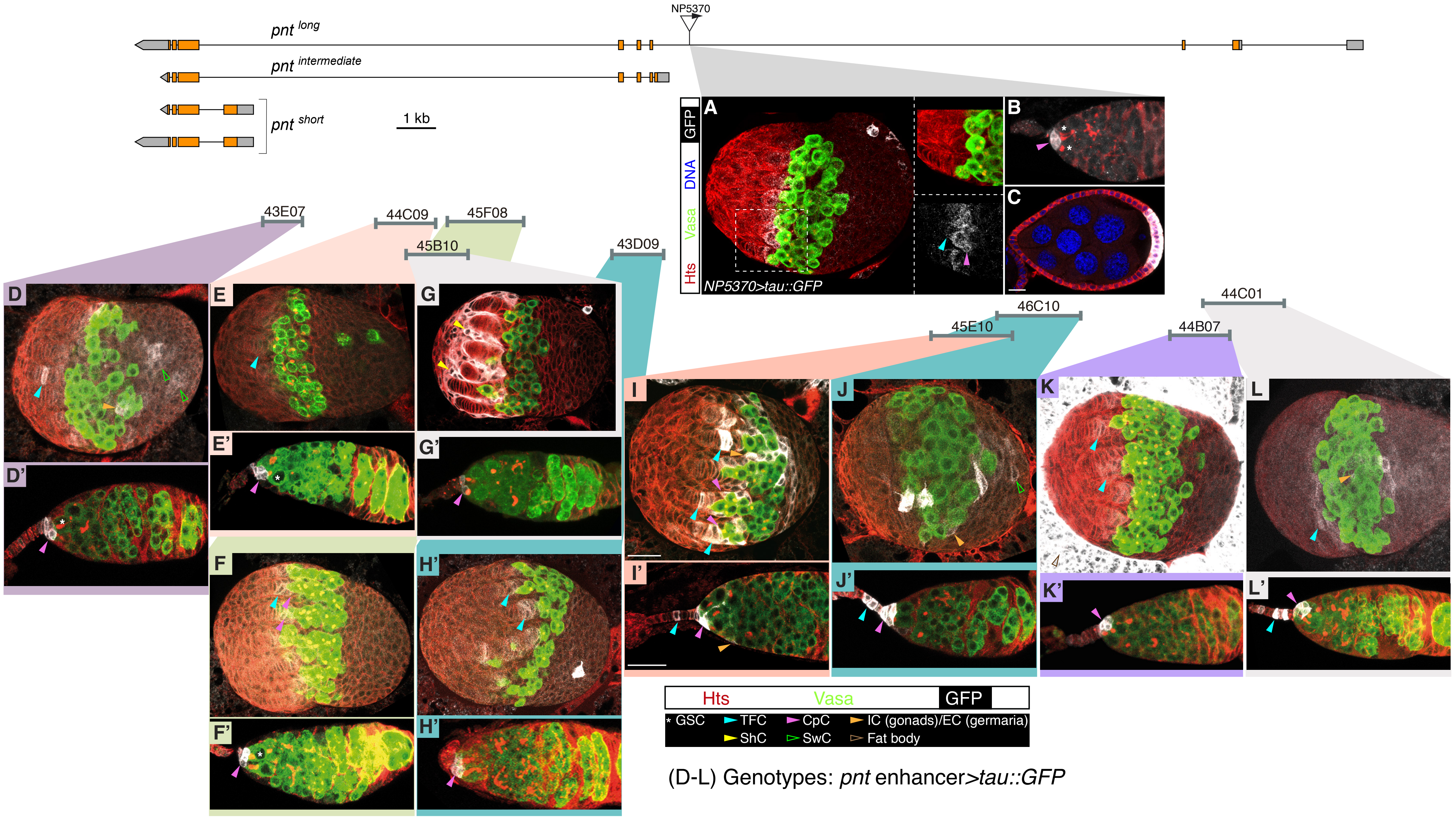
NP5370
pnt long
pnt intermediate
pnt short
1 kb
GFP
DNA
Vasa
Hts
NP5370>tau::GFP
A
B
C
43E07
44C09
45F08
45B10
43D09
46C10
45E10
44C01
44B07
D
D'
E
E'
F
F'
G
G'
H'
H'
I
I'
J
J'
K
K'
L
L'
Hts Vasa GFP
* GSC
TFC
CpC
IC (gonads)/EC (germaria)
ShC
SwC
Fat body
(D-L) Genotypes: pnt enhancer>tau::GFP

**Figure S1**

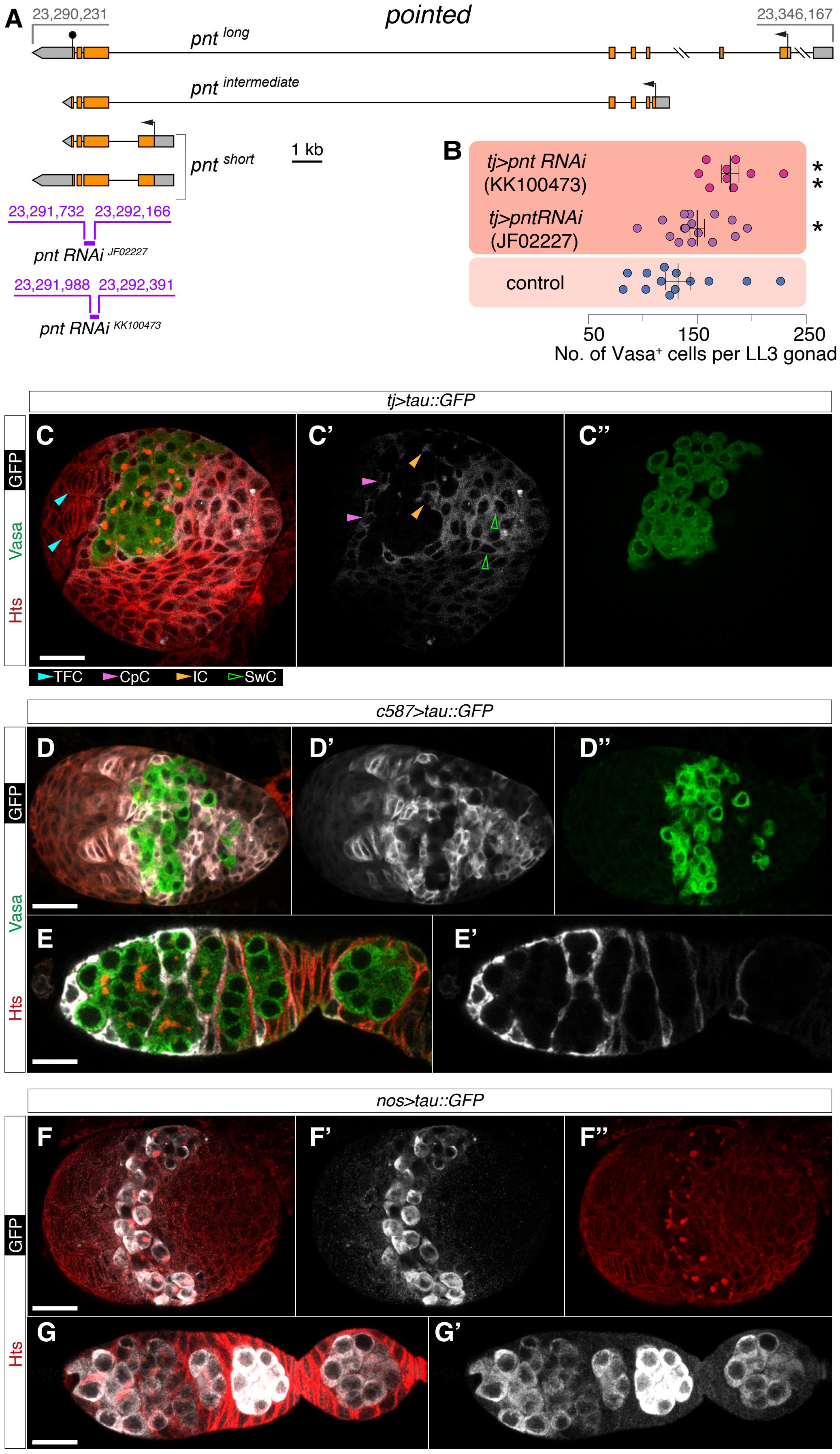

A
pointed
23,290,231
23,346,167
pnt long
pnt intermediate
pnt short
1 kb
23,291,732 23,292,166
pnt RNAi JF02227
23,291,988 23,292,391
pnt RNAi KK100473
B
tj>pnt RNAi (KK100473)
tj>pntRNAi (JF02227)
control
*
*
*
50
150
250
No. of Vasa+ cells per LL3 gonad
tj>tau::GFP
C
C'
C''
GFP
Vasa
Hts
TFC
CpC
IC
SwC
c587>tau::GFP
D
D'
D''
E
E'
nos>tau::GFP
F
F'
F''
G
G'


**Figure S2**

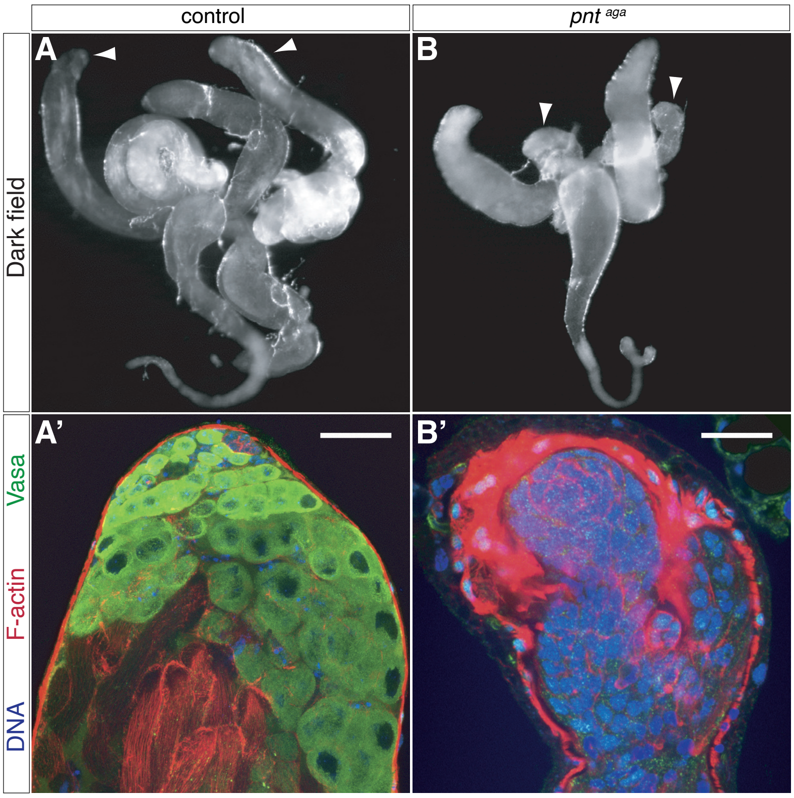


**Figure S3**

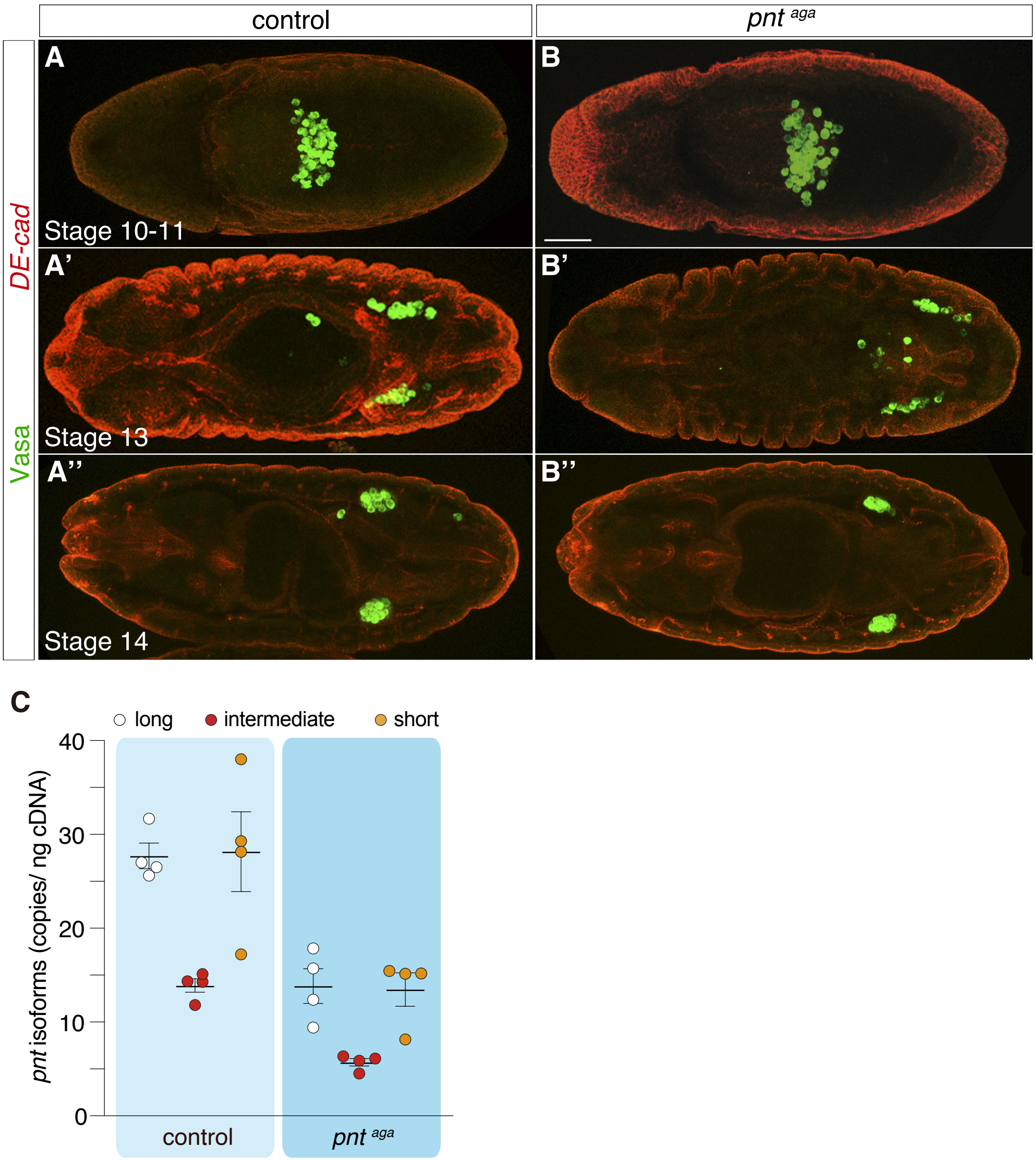
control
pnt aga
A
B
DE-cad
Vasa
Stage 10-11
A'
B'
Stage 13
A''
B''
Stage 14
C
long
intermediate
short
pnt isoforms (copies/ ng cDNA)
40
30
20
10
0
control
pnt aga

**Figure S4**

**A**

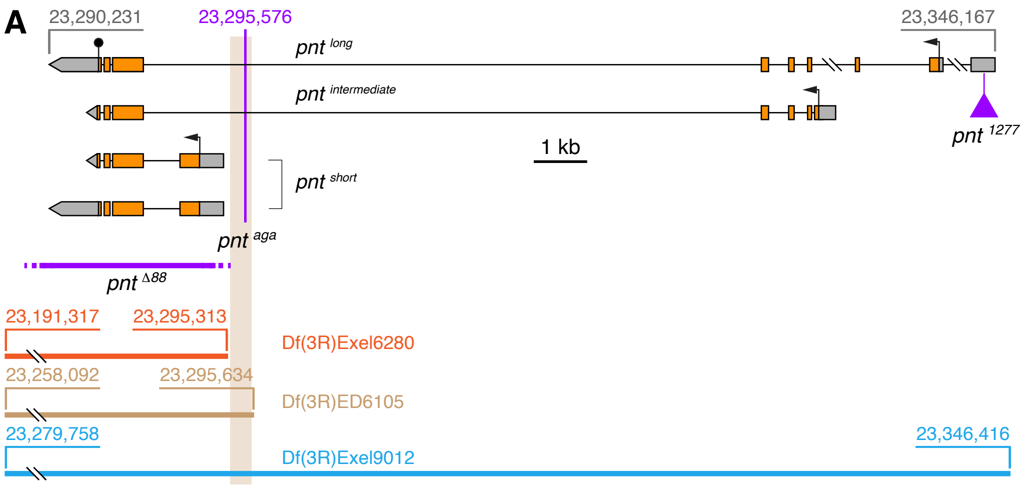


**B**

Sequence: *white*⁻ control (query) vs. reference Flybase genome (sbjct)

```
Query  290         CCGTGTGTGGGAGAGAGCGCTTCGTGTGTACTTCGGCTCGACGACTTCGACTCGCATTCC    350
                   ||||||||||||||||||||||||||||||||||||||||||||||||||||||||||||
Sbjct  23,295,526  CCGTGTGTGGGAGAGAGCGCTTCGTGTGTACTTCGGCTCGACGACTTCGACTCGCATTCC    23,295,586
```

Sequence: mutant sequence (query) vs. reference Flybase genome (sbjct)

```
Query  290         CCGTGTGTGGGAGAGAGCGCTTCGTGTGTACTTCGGCTCGACGACTTCG-CTTCGCATTC    348
                   |||||||||||||||||||||||||||||||||||||||||||||||||  | ||||||||
Sbjct  23,295,526  CCGTGTGTGGGAGAGAGCGCTTCGTGTGTACTTCGGCTCGACGACTTCGAC-TCGCATTC    23,295,585
```

**C**

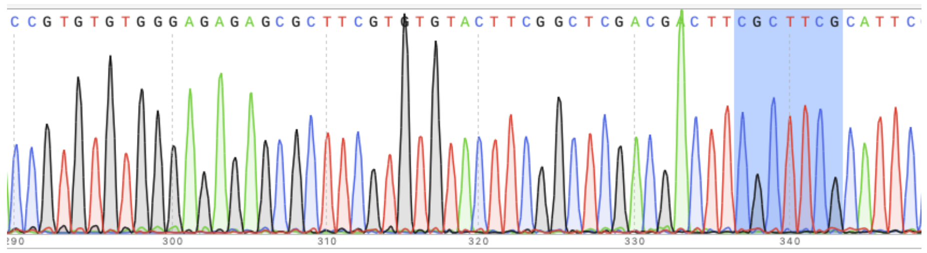


**D**

| Genotype | Viability | Phenotype |
|---|---|---|
| *pnt* Δ88/ Df(3R)Exel6280 | lethal | - - - |
| *pnt* Δ88/ Df(3R)Exel9012 | lethal | - - - |
| *pnt* Δ88/ Df(3R)ED6105 | lethal | - - - |
| *pnt* aga/ *pnt* Δ88 | viable | normal ovaries |
| *pnt* aga/ *pnt* 1277 | viable | normal ovaries |
| *pnt* aga/ Df(3R)Exel6280 | viable | normal ovaries |
| *pnt* aga/ Df(3R)ED6105 | viable | agametic ovaries |
| *pnt* aga/ Df(3R)Exel9012 | viable | agametic ovaries |

**Figure S5**

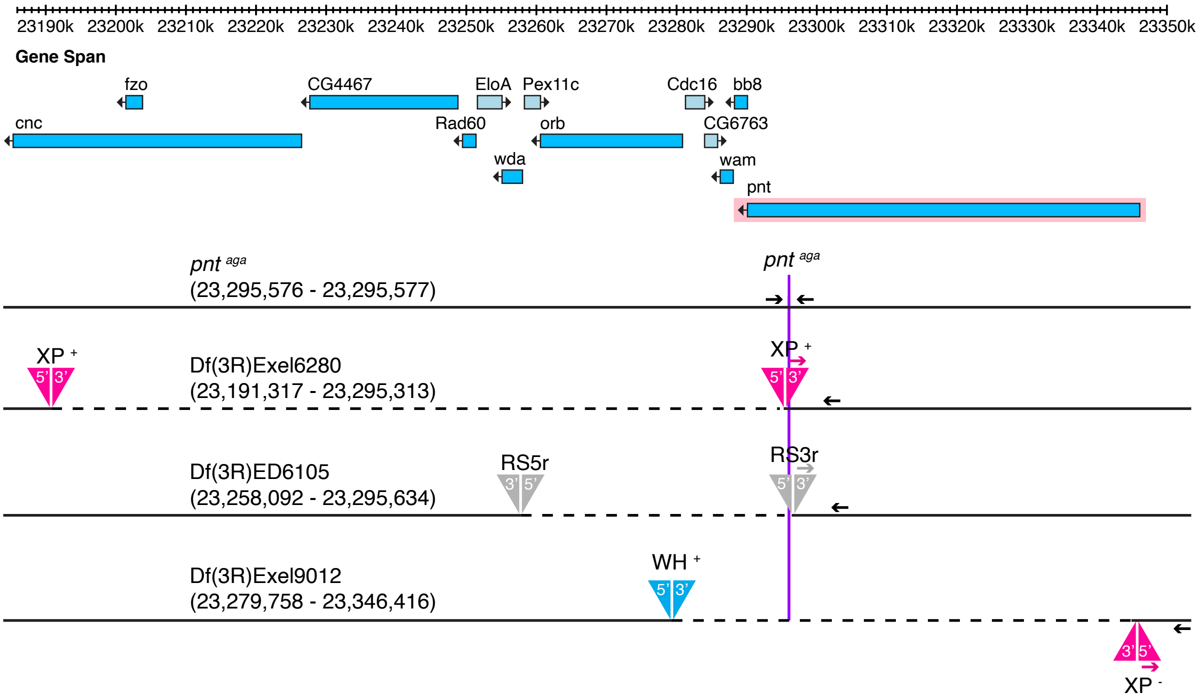
23190k 23200k 23210k 23220k 23230k 23240k 23250k 23260k 23270k 23280k 23290k 23300k 23310k 23320k 23330k 23340k 23350k
Gene Span
fzo
CG4467
EloA
Pex11c
Cdc16
bb8
cnc
Rad60
orb
CG6763
wda
wam
pnt
pnt aga
(23,295,576 - 23,295,577)
pnt aga
XP +
Df(3R)Exel6280
(23,191,317 - 23,295,313)
XP +
Df(3R)ED6105
(23,258,092 - 23,295,634)
RS5r
RS3r
Df(3R)Exel9012
(23,279,758 - 23,346,416)
WH +
XP -

**Figure S6**
